# A post-mitotic bistable switch actively safeguards mature neurons against reciprocal transdifferentiation *in vivo*

**DOI:** 10.64898/2026.06.19.733455

**Authors:** Qi-Nan Chen, Yineng Xu, Bei Wang, Yifan Shen, Sasidhar Karuparti, Nambita S. Sahai, Chun Han

## Abstract

How mature cells actively defend their identity against alternative fates is poorly understood. Here we show that two types of fully differentiated *Drosophila* somatosensory neurons, derived from distinct lineages, are continuously prevented from transdifferentiating into one another by a post-mitotic bistable switch. Two homeodomain transcription factors, Hmx and Cut, are co-expressed in both neuronal types but adopt opposite stoichiometric dominances through mutual repression, each defining one identity. Disrupting this balance in mature neurons triggers deterministic, lineage-crossing reprogramming of transcriptome, dendritic and axonal architecture, and sensory behavior without reversion to a progenitor state. This regulatory logic is evolutionarily conserved: *Drosophila* sensory identities share transcriptomic signatures with distinct human Aδ mechanoreceptors, and human CUX and HMX proteins functionally replace their fly counterparts. Mature neuronal identity is thus not a fixed endpoint but a poised equilibrium actively maintained by ongoing competition between opposing selectors.

**TEASER:** Mature somatosensory identity is a poised equilibrium actively maintained by continuous competition between opposing terminal selectors.

## INTRODUCTION

A fundamental property of complex multicellular organisms is the stability of cell identity. This stability is particularly important for the functional complexity of the nervous system, in which diverse neuronal subtypes must reliably execute specialized roles over an organism’s lifespan. Classically, terminal differentiation is viewed as a one-way trajectory in which cell identity, once acquired, is locked in by restrictive chromatin landscapes and stable transcriptional programs. Yet the discovery that “terminal selector” transcription factors must be continuously expressed in mature neurons to sustain subtype-specific features (*1–3*) has revealed that even fully differentiated identities are actively maintained rather than passively inherited. This raises a fundamental question regarding cellular plasticity and circuit fidelity: do mature neurons simply drift toward a degraded or generic state in the absence of maintenance factors, or do they actively suppress specific competing identities to prevent the cells from adopting an alternative fate? The latter possibility would imply that a differentiated cell is held in a poised, metastable equilibrium, continuously prevented from transdifferentiating into a competing identity.

In the prevailing view of identity maintenance, terminal selectors lock in cell fate primarily through positive mechanisms: they directly activate batteries of subtype-specific effector genes and sustain their own expression via autoregulatory feedback loops (*2, 4*). Some terminal selectors additionally repress effector genes of alternative, closely related neuronal identities (*2, 5, 6*). However, it remains unclear whether such repression merely buffers transcriptional noise within a lineage, or instead constitutes a primary, instructive force that defines the discreteness of cellular identity. Furthermore, while terminal selectors are known to prevent identity drifts within the same lineages (*3, 7*), it is unknown if similar logic operates to prevent transdifferentiation across distinct progenitor lineages, and whether such regulatory circuits are conserved across evolution.

The *Drosophila* dendritic arborization (da) sensory neurons offer a uniquely tractable system to dissect these questions *in vivo*. Innervating the larval body wall, da neurons are divided into four morphologically and functionally distinct classes (C1da–C4da) (*8, 9*). Critically, unlike many neuronal subtypes that arise from the asymmetric division of a single mother cell, each da class is generated from a separate progenitor lineage (*10*), making them an ideal system to ask whether terminal identity is actively defended across deep lineage boundaries. Despite these independent developmental origins, neighboring proneural clusters co-express overlapping batteries of transcription factors (*9*), creating a paradox: how do lineages with partially shared transcriptional programs resolve into permanently discrete, non-interchangeable identities? This challenge is especially clear between Class II (C2da) and Class III (C3da) neurons, which arise from distinct lineages yet share transcription factor expression and innervate the same epidermal territory, while exhibiting strikingly different dendritic architectures and sensory functions: C2da neurons elaborate sparse arbors decorated with terminal varicosities and mediate proprioceptive feedback during locomotion (*8, 11*), whereas C3da neurons elaborate highly branched arbors decorated with characteristic dendritic spikes and mediate gentle touch and noxious cold sensation (*12, 13*). While the homeodomain protein Cut (Ct) is known to correlate with dendrite complexity (*14*), the instructive logic governing these precise identity boundaries has remained elusive.

Here, we uncover a post-mitotic bistable switch that establishes and actively defends the identities of C2da and C3da somatosensory lineages. We show that the homeodomain transcription factors H6-like-homeobox (Hmx) and Ct function as opposing terminal selectors for C2da and C3da neurons, respectively. Although Hmx and Ct are co-expressed in both lineages from the progenitor stage onward, they establish opposite stoichiometric dominances in mature C2da and C3da neurons. Importantly, maintenance of neuronal identity requires active, post-mitotic mutual repression: Ct actively suppresses the C2da selector Hmx to generate the complex C3da phenotype, while Hmx protects the basal C2da-like state by repressing Ct. Strikingly, disrupting this asymmetric dominance in mature neurons triggers deterministic, lineage-crossing transdifferentiation, reprogramming the entire cellular phenotype to the opposite neuronal identity. Finally, the molecular components of this switch are evolutionarily conserved: *Drosophila* C2da and C3da neurons share transcriptomic signatures with human Aδ high- and low-threshold mechanoreceptors, respectively, and human CUX and HMX proteins functionally substitute for their fly counterparts to drive identity interconversion *in vivo*. Together, these findings reveal that discrete cellular identities are not passively inherited but are sustained throughout life by an active, mutually antagonistic competition between opposing terminal selectors, a logic that may be broadly deployed to safeguard cell fate across diverse lineages and species.

## RESULTS

### Cut and Hmx act as mutually antagonistic terminal selectors for two somatosensory lineages

To understand how neuronal identities of da neurons are maintained, we established a system to simultaneously visualize multiple subtypes in live animals. We identified the FlyLight (*15*) *R80C08* enhancer as a specific marker for C1da and C2da neurons. Membrane-targeted tdTomato (CD4-tdTom) (*16*) fused to *R80C08* (*R80C08-CD4-tdTom*) labels all larval C1da and C2da neurons (Figure 1A, showing dorsal C1da neurons ddaD and ddaE and the C2da neuron ddaB; Supplementary Table 1). We similarly constructed *R20C11-CD4-tdGFP*, which labels a subset of C2da neurons, including ddaB (Figure S1A; Supplementary Table 1). Conversely, we labeled C3da neurons using the C3da-specific *LexA^nompC^*(*17*), driving *LexAOP2-CD4-IFP2.0-2A-HO1* (*CD4-IFP*), or a C3da-specific *ss* enhancer (*18*) driving *CD4-tdTom* (Figure 1A and S1A, showing dorsal C3da neurons ddaA and ddaF; Supplementary Table 1). Using the pan-da Gal4 drivers *Gal4^21-7^* (*19*) and *Gal4^RluA1^* (*20*) we manipulated gene expression in all da neurons post-mitotically and examined the consequences in wandering 3^rd^ instar larvae.

**Figure 1.**
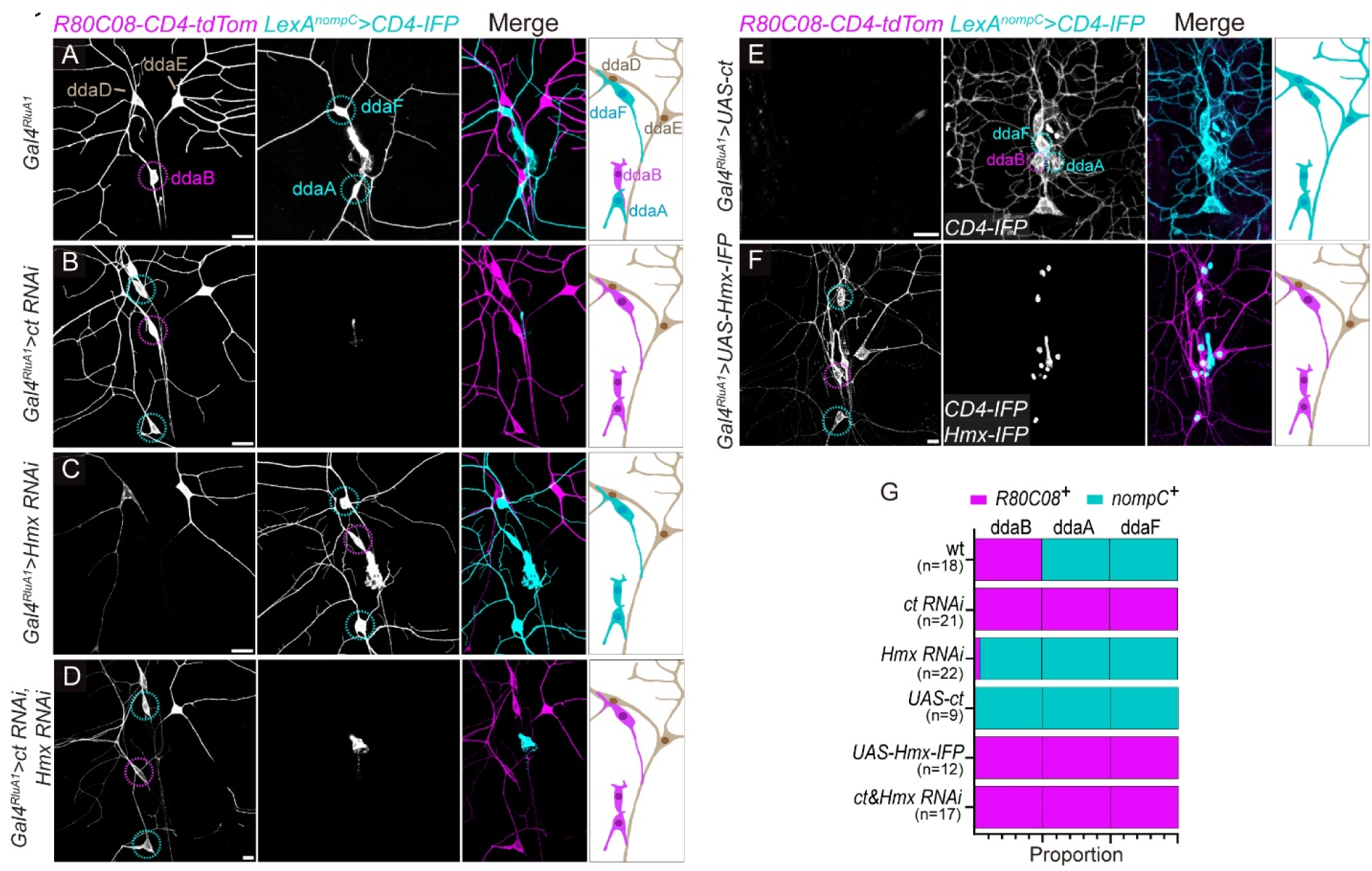
Cut and Hmx act as mutually antagonistic terminal selectors for two somatosensory lineages. **(A-F)** Expression of markers (*R80C08-CD4-tdTom* for C1da/C2da and *LexA^nompC^>CD4-IFP* for C3da) in dorsal da neurons of WT control (**A**), *ct RNAi* (**B**), *Hmx RNAi* (**C**), *ct* and *Hmx* double knockdown (KD) (**D**), *ct* overexpression (OE) (*UAS-ct*) (**E**), and *Hmx* OE (*UAS-Hmx-IFP*) (**F**). The pan-da *Gal4^RluA1^* drives transgene expression. Larvae were examined at the late 3rd instar, except for *UAS-ct* (**E**), which were examined at the 1st instar due to early larval lethality. Cyan circles indicate C3da neurons (ddaA/F); magenta circles indicate C2da neurons (ddaB). The diagrams on the right illustrate cell conversion outcomes. **(G)** Neuron proportions based on marker expression in **(A–F)**. Sample sizes (n = number of neurons, N = number of animals): Control (n = 18, N = 8); *ct RNAi* (n = 21, N = 6); *Hmx RNAi* (n = 22, N = 11); *UAS-ct* (n = 9, N = 6); *UAS-Hmx-IFP* (n = 12, N = 6); and *ct* & *Hmx* RNAi (n = 17, N= 10). Scale bars: 20 μm in (**A–C**); 10 μm in (**D-F**).

Ct expression levels are known to correlate with dendrite complexity of da neurons, with high Ct levels in elaborated C3da and C4da neurons, low level in simpler C2da neurons, and no expression in C1da neurons (*14*). To determine if Ct also regulates other class-specific properties, we knocked it down (KD) in all da neurons. Strikingly, *ct* KD did not merely simplify C3da dendrites as previously reported (*14*), but caused a complete switch of class-specific marker expression: ddaA and ddaF lost the C3da marker *LexA^nompC^>CD4-IFP* and ectopically expressed the C1da/C2da marker *R80C08-CD4-tdTom* (Figure 1B). This switch was corroborated by reductions of *ss-CD4-tdTom* in all C3da neurons and gain of *R20C11-CD4-tdGFP* in ddaF (Figure S1B). In these animals, marker expression was unaltered in other da classes, suggesting that *ct* loss does not generally disrupt cell identity but causes a specific C3da-to-C2da identity conversion.

Since *R80C08* is derived from the *H6-like-homeobox* (*Hmx*) locus, we hypothesized that the homeodomain protein Hmx is related to C1da/C2da identity. Interestingly, *Hmx* KD in all da neurons caused a reciprocal switch of marker expression in C2da neurons only: ddaB lost *R80C08-CD4-tdTom* and *R20C11-CD4-tdGFP* expression while gaining C3da markers *LexA^nompC^>CD4-IFP* and *ss-CD4-tdTom* (Figures 1C and S1C), suggesting a C2da-to-C3da conversion. These marker switches were highly penetrant (100% for *ct* KD in ddaA/F neurons; 90% for Hmx KD in ddaB; Figure 1G). Notably, while all C3da neurons switched upon *ct* KD, only specific C2da neurons (ddaB and vdaC) switched upon *Hmx* KD (Supplementary Table 1), suggesting context-dependent sensitivity or redundancy in other C2da neurons (IdaA and vdaA).

Given that *ct* and *Hmx* are required in two distinct, apparently opposing neuronal subtypes to prevent reciprocal marker conversions, we asked if forcing Ct and Hmx expression in the opposite cell type can override the endogenous program. Indeed, pan-da overexpression (OE) of Ct converted C2da neurons (ddaB) into C3da-like cells, as evidenced by the loss of *R80C08-CD4-tdTom* and gain *LexA^nompC^>CD4-IFP* (Figure 1E). Interestingly, C1da (ddaD/E) neurons showed a similar switch of marker expression as ddaB (Figure 1E). Reciprocally, Hmx OE converted C3da neurons (ddaA/F) into C2da-like cells, indicated by marker expression (Figure 1F). These switches were 100% penetrant (Figure 1G).

The above results suggest that C2da and C3da are binary identity choices for both lineages, and that Ct and Hmx act as mutually antagonistic terminal selectors to lock neurons into one of these identities. To determine the hierarchy of the two identities, we performed a double knockdown. In *ct/Hmx* double-KD larvae, ddaA, ddaF, and ddaB all expressed the C2da marker *R80C08-CD4-tdTom* but not the C3da marker *LexA^nompC^>CD4-IFP* (Figures 1D and 1G), suggesting that C2da is the default fate for both lineages in the absence of Ct and Hmx.

### Ct and Hmx manipulations reprogram class-specific dendritic morphogenesis

We next assessed whether the reprogramming extended to dendritic patterning, a definitive feature of neuronal identity. C2da neurons show simpler branches with terminal swellings, or “varicosities” (Figure 2A), whereas C3da neurons display more complex arbors decorated with numerous short terminal branches called dendritic spikes (Figure 2D and 2G) (*8*). Using the *CoinFlp* system (*21*) to generate sparse, single-cell clones expressing RNAi or wildtype (WT) proteins (Figures S2A-S2G), we quantified dendritic architecture. With the KD approach, *Hmx*-deficient ddaB clones acquired C3da-like complexity, exhibiting a 143% increase in terminal branches, a 25% increase in dendrite length, and the emergence of dendritic spikes with simultaneous loss of varicosities (Figures 2B, 2J, and 2K). Conversely, *ct*-deficient ddaF and ddaA clones lost the C3da morphology and adopted a simplified, C2da-like architecture, characterized by ddaB-level terminal branch numbers and total dendrite length, as well as the appearance of distal varicosity (Figures 2E, 2H, 2J, and 2K). Similarly, the OE approaches also resulted in reprograming of dendritic receptive fields: Ct OE in ddaB clones induced a robust C3da phenotype, with a 136% increase in terminal branches and 42% increase in length, along with loss of varicosities and emergence of dendritic spikes (Figures 2C, 2J, and 2K). In contrast, Hmx OE in ddaA/F clones simplified the dendritic arbor, strongly reducing branch number and length (Figures 2F, 2I, 2J and 2K). While their dendrites were not reduced to the ddaB level, these neurons showed C2da-like varicosities, suggesting substantial, if not complete, dendritic transformation.

**Figure 2.**
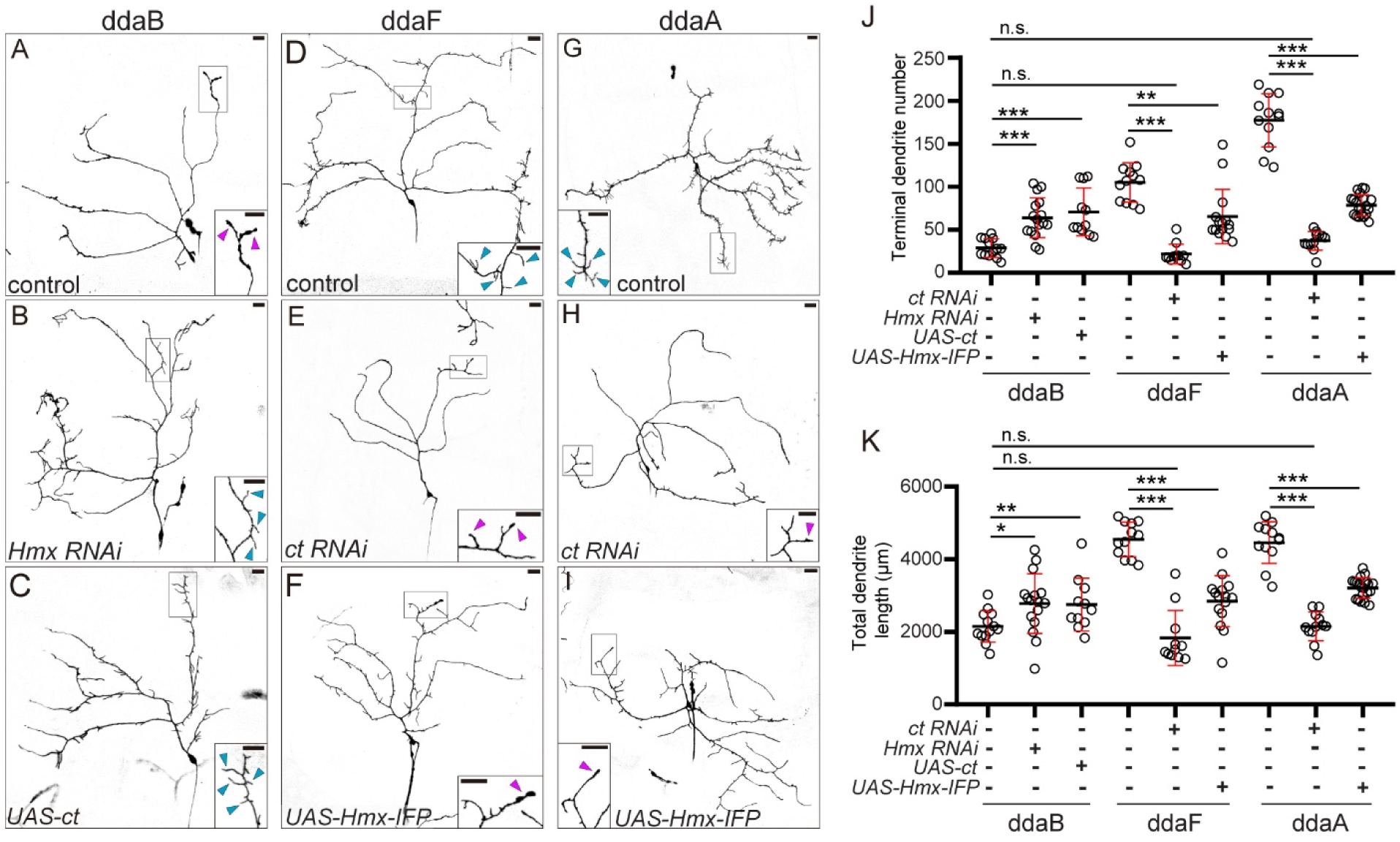
Ct and Hmx manipulations reprogram class-specific dendritic morphogenesis. **(A–I)** CoinFlp clones of ddaB (C2da) (**A-C**) and C3da ddaF (**D-F**) and ddaA neurons (G-I) in control (**A, D, G**), *Hmx* RNAi (**B**), *UAS-ct* (**C**), *ct* RNAi (**E, H**), and *UAS-Hmx-IFP* (**F**, **l**). Insets show higher-magnification views of dendritic terminals. Cyan arrowheads indicate dendritic spikes (characteristic protrusions of C3da); magenta arrowheads indicate varicosities (characteristic of C2da). **(J and K)** Terminal dendrite number (**J**) and total dendrite length (**K**) across indicated genotypes. Sample sizes (n = neurons, N = animals): WT ddaB (n = 13, N = 11), *Hmx RNAi* ddaB (n = 16, N = 12); *UAS-ct* ddaB (n = 11, N = 11); WT ddaF (n = 12, N = 10), *ct RNAi* ddaF (n = 11, N = 10); *UAS-Hmx-IFP* ddaF (n = 15, N = 11); WT ddaA (n = 12, N = 10), *ct RNAi* ddaA (n = 12, N = 10); *UAS-Hmx-IFP* ddaA (n = 20, N = 18). One-way ANOVA with Tukey post-hoc (*p<0.05, **p<0.01, ***p<0.001, n.s., not significant). Black bars, mean; red bars, SD. Scale bars: 25 μm.

### Ct and Hmx control the entire transcriptional identity of C2da and C3da neurons

To assess the molecular completeness of these C2da/C3da interconversions, we performed single-nucleus RNA sequencing (snRNA-seq) on ∼15,000 larval body-wall cells from WT control (8953 cells), *ct* KD (6532 cells), and *Hmx* KD (6152 cells) genotypes (Figure S3A). Nuclei of da neurons were labeled by *Gal4^RluA1^>Lamin-EGFP* to facilitate nuclear sorting and subsequent analysis. Based on expressions of EGFP, *ct*, and *Hmx*, we identified a “sensory neuron” cluster that should encompass all da neurons (Figure S3B). We further identified subclusters that correspond to the four da subtypes based on class-specific markers (*nompC* for C3da; *ppk* for C4da; *ab* for C1da) (*12, 22–25*) (Figures S3C and S3D).

*ct* KD resulted in a massive shift in population structure: the proportion of identified C2da neurons expanded (12.2% to 23.3%), while C3da neurons collapsed (24.0% to 7.6%) (Figures 3A and 3B). This confirms that prespecified C3da neurons shifted their transcriptomic profile to align with C2da neurons. Reciprocally, *Hmx* KD expanded the C3da population (24.0% to 29.9%) at the expense of C2da neurons (12.2% to 10.5%) (Figures 3A and 3B). *Hmx* KD caused smaller effect sizes than *ct* KD, consistent with only two of the four C2da neurons showing marker switches (Supplementary Table 1). *ct* and *Hmx* KDs also caused small reductions of C1da (1.5% and 2.2%, respectively) and C4da (3.9% and 2.5%, respectively) neurons (Figures 3A and 3B), which could be due to partial transcriptomic alterations in these cells.

**Figure 3.**
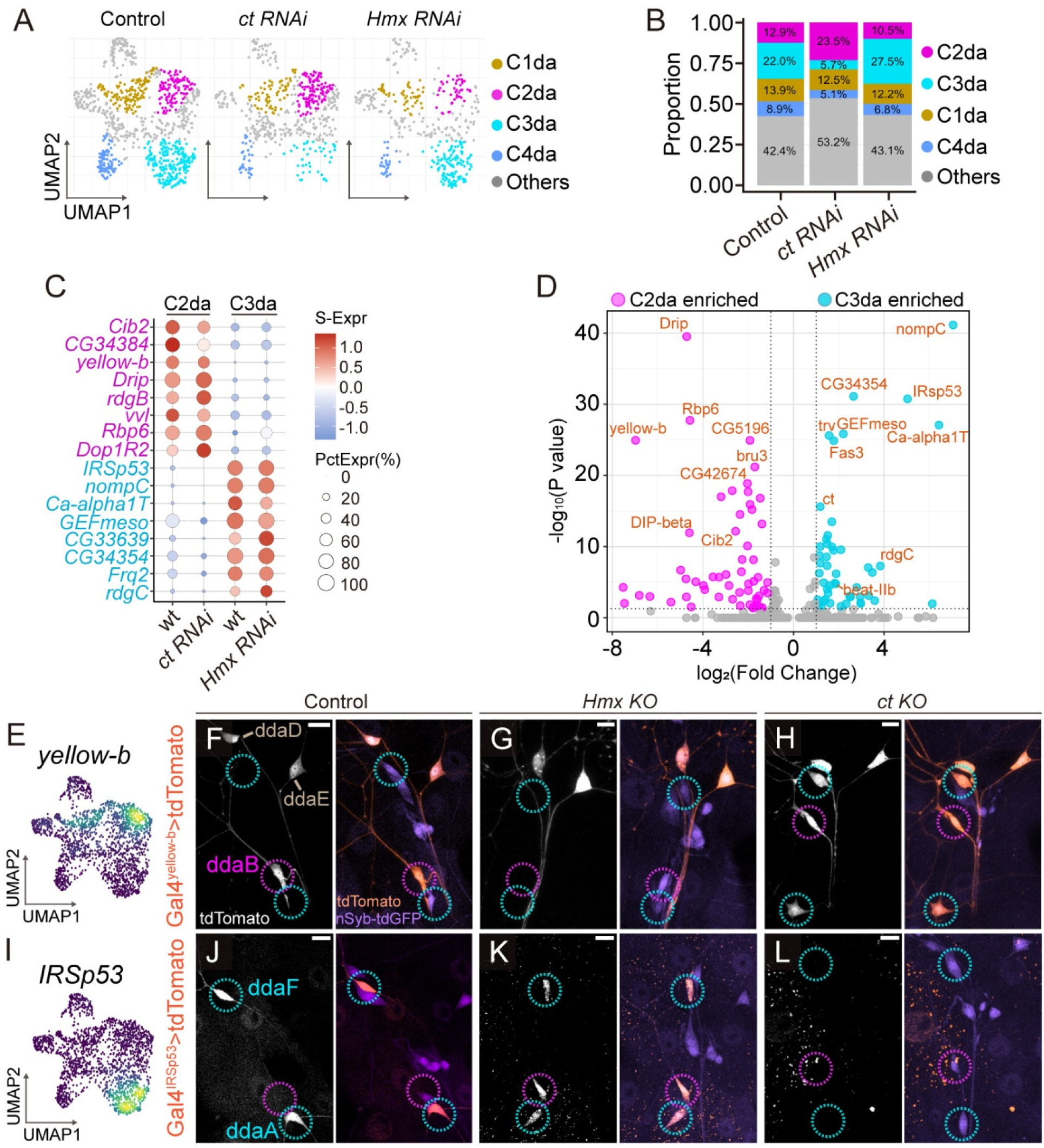
Ct and Hmx control the entire transcriptional identity of C2da and C3da neurons. **(A)** UMAP of the sensory neuron cluster in control, *ct RNAi*, and *Hmx RNAi* datasets, with putative da subtypes labeled in different colors. **(B)** Proportions of da classes in (**A**). Percentages are relative to the total cell number in each genotype. **(C)** Expression profiles of C2da- and C3da-enriched genes in C2da and C3da cell clusters. Dot color: scaled average expression (S-Expr); dot size: percentage of expressing cells (PctExpr). **(D)** Volcano plot showing differentially expressed genes (DEGs) between wild-type C2da and C3da neurons. A subset of the DEGs are labeled. **(E-L)** Validation of differentially expressed genes (DEGs) *yellow-b* (C2da-specific, **F-H**) and *IRSp53* (C3da-specific, **J-L**) with corresponding Gal4 reporters. UMAP plots (**E, I**) serve as single-cell expression references. The respective Gal4 lines drive *UAS-tdTomato* expression in Control (**F, J**), *Hmx KO* (**G, K**), and *ct KO* (**H, L**) larvae. The pan-neuronal marker *nSyb-tdGFP* was included to facilitate identification of neurons. Magenta circles indicate C2da (ddaB) neurons, and cyan circles indicate C3da (ddaA/ddaF) neurons. Scale bars: 10 μm.

Differential expression (DE) analysis of WT cells identified genes enriched in each da class (Figures 3C, 3D, S3E). Crucially, the DE genes (DEGs) driving these class distinctions included specific ion channels, receptors, and synaptic adhesion molecules which are likely required to execute subtype-specific sensory functions. C2da neurons in *ct* KD animals expressed the DEGs at levels indistinguishable from wild-type C2da neurons (Figures 3C and S3E). Interestingly, the few remaining C3da neurons in *ct* KD animals showed dampened expression of C3da genes, suggesting a “partial” loss of identity in cells that had not yet fully switched. Similarly, C3da neurons in *Hmx* KD are indistinguishable from wild-type C3da neurons (Figures 3C and S3E). These data suggest that the “converted” neurons were transcriptomically homologous to their target fates.

To validate these transcriptomic changes *in vivo* using an RNAi-independent approach, we obtained or generated Gal4 reporters for a subset of identified DE genes (C2da: *yellow-b*, *Rbp6*; C3da: *rdgC*, *IRSp53*, *beat-IIb*, *Fas3*) (Figures 3D, 3E, 3I, S4) and verified their class-specific expression patterns in WT neurons (Figures 3F, 3J, S4). We then knocked out (KO) *ct* and *Hmx* in da lineages using tissue-specific Cas9. We generated *asense* (*ase*)*-Cas9*, active in committed progenitors of sensory neurons (*26*), to avoid perdurance (gene products made before mutagenesis masking LOF phenotypes) associated with purely post-mitotic mutagenesis (*27*). *ase-Cas9* combined with guide RNAs (gRNAs) targeting *ct* and *Hmx* produced the expected class-switching changes of *Gal4^R80C08^* and *Gal4^nompC^*expression in da neurons (Figures S4A-S4F), validating the effectiveness of this approach. Furthermore, *Hmx* KO abolished C2da reporters and ectopically induced C3da reporters in ddaB, while *ct* KO reciprocally silenced C3da reporters and induced C2da reporters in ddaA/F (Figures 3G, 3H, 3K, 3L, S4). These data confirm that *Ct* and *Hmx* function as mutually antagonizing master regulators, directly or indirectly controlling the entire battery of subtype-specific effector genes defining C2da and C3da identities.

### Ct and Hmx in progenitors seed asymmetric dominance in post-mitotic neurons through mutual repression

Our transcriptomic data show that Hmx and Ct are expressed in both C2da and C3da neurons (Figure S3D), even though they each specify only one lineage. To resolve this paradox, we investigated the pre-mitotic roles of *Hmx* and *ct* by knocking them out using *SOP-Cas9* (*27*), which is driven by the *scute* (*sc*) enhancer that is already active in proneural precursors (*28*). We found that early KO of either *ct* or *Hmx* led to the loss of both C2da and C3da neurons (Figures 4A-4F), indicating that both transcription factors are co-expressed in precursors and functionally required for early survival or specification across these distinct lineages. Importantly, this early co-requirement is highly asymmetric: *ct* KO caused most frequent loss of C3da neurons but also affected C2da neurons to a lesser degree (Figures 4C and 4D), while *Hmx* KO predominantly eliminated C2da neurons but also caused a milder loss of C3da neurons (Figures 4E and 4F). This asymmetric co-requirement suggests that Cut and Hmx are co-expressed early in both cell lineages, but they each plays a dominant role in the progenitors of the neuronal subtypes they maintain.

**Figure 4.**
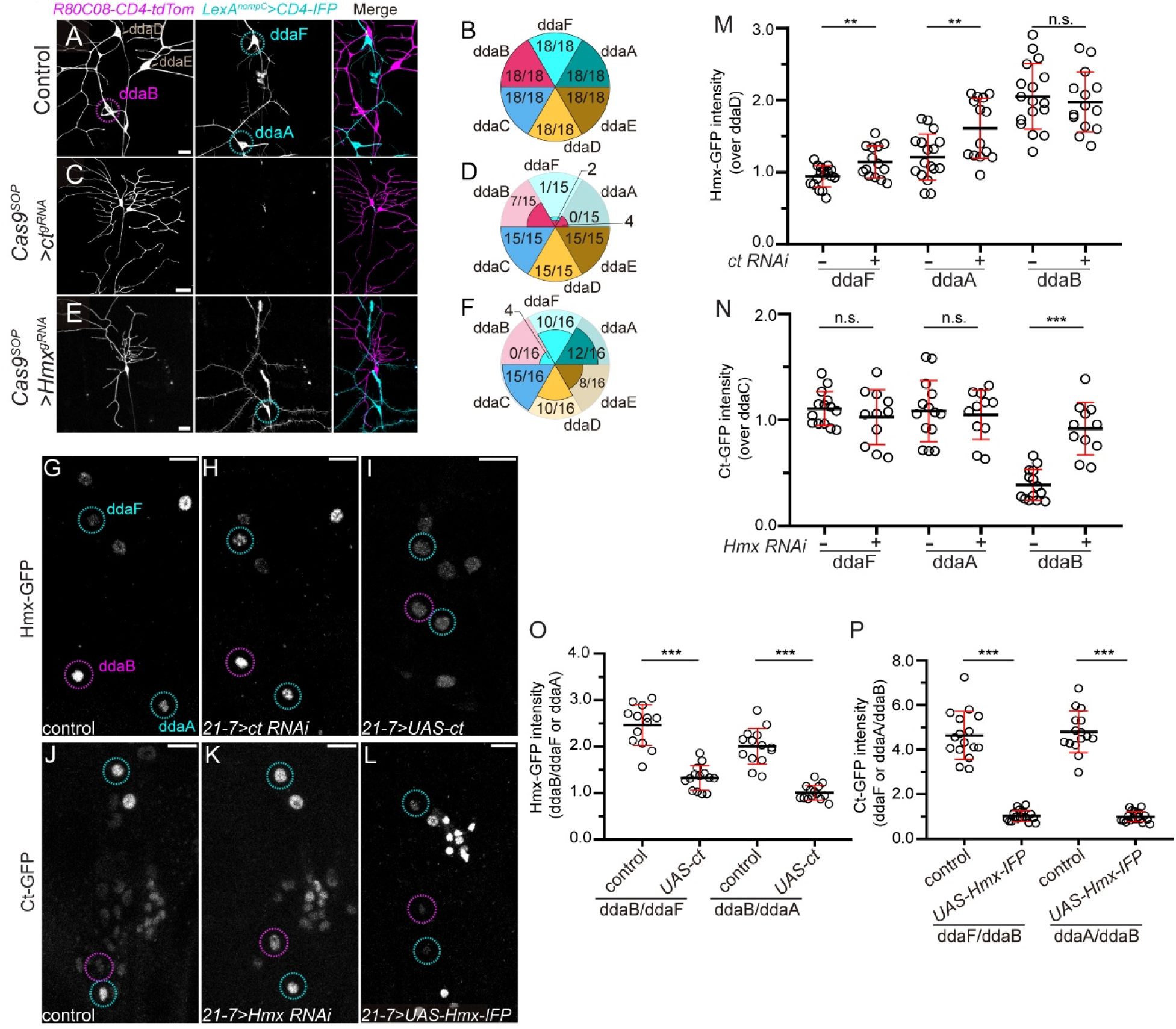
Ct and Hmx in progenitors seed asymmetric dominance in post-mitotic neurons through mutual repression. **(A-E)** Marker expression and da neuron counting in WT control (**A, B**) and after pre-mitotic KO of *ct* (**C, D**) and *Hmx* (**E, F**) by *SOP-Cas9*. Cyan circles indicate C3da neurons (ddaF/A); magenta circles indicate C2da neurons (ddaB). In the pie charts, each sector represents a da neuron in the dorsal da cluster. The fraction inside a sector indicates the survival rate (observed cells with the correct identity / total hemi-segments examined). Neurons exhibiting marker conversion are represented with colors matching the converted identities, and their numbers are shown outside the sector. **(G-I)** Endogenous *Hmx-GFP* expression in late 3^rd^ instar larvae of the WT control (**G**) and *ct* KD (**H**) and *ct* OE (*UAS-ct*) (**I**). **(J-L)** Endogenous *ct-GFP* expression in late 3rd instar larvae of Control (**J**), *Hmx RNAi* (**K**), and *Hmx* OE (*UAS-Hmx-IFP*) (**L**). All KD and OE were driven by *Gal4^21-7^*. Identification of neurons was facilitated by *UAS-CD4-IFP* expression (not shown). Cyan circles indicate C3da neurons (ddaF/A); magenta circles indicate C2da neurons (ddaB). **(M)** Hmx-GFP intensities in C2da (ddaB) and C3da (ddaA/F) neurons of WT and *ct* RNAi, normalized by those in C1da (ddaD) neurons. (**N**) Ct-GFP intensities in C2da (ddaB) and C3da (ddaA/F) neurons of WT and *Hmx* RNAi, normalized by those in C4da (ddaC) neurons. (**O**) Hmx-GFP intensities in C2da (ddaB) neurons of *ct* OE, normalized by those in C3da (ddaA/F) neurons. **(P)** Ct-GFP intensities in C3da (ddaA/F) of *Hmx* OE, normalized by those in C2da (ddaB) neurons. Sample sizes (n = neurons): n = 12–18 per genotype. One-way ANOVA with Tukey post-hoc (**p<0.01, ***p<0.001; n.s., not significant). Black bars, mean; red bars, SD. In all image panels, scale bars, 10 μm.

To understand how Hmx and Ct antagonize each other in C2da and C3da neurons at the protein level, we tagged endogenous Hmx and Ct proteins with GFP at their C-termini through CRISPR-mediated knock-in (KI). Hmx-GFP and Ct-GFP signals were detected in the nuclei of da neurons and were eliminated upon *Hmx* and *ct* KD, respectively (Figures S5A-S5J), cross-validating both KI alleles and the effectiveness of KD.

In WT larvae, we observed a high Hmx-GFP level in the C2da neuron ddaB and much lower levels in C3da ddaA (41% less) and ddaF (54% less) (Figures 4G and 4M). *ct* KD appreciably elevated Hmx-GFP in ddaA (33% increase) and ddaF (21% increase) without affecting it in ddaB (Figures 4H and 4M). In contrast, Ct overexpression reduced Hmx-GFP in ddaB to a level comparable to those in ddaA and ddaF (Figures 4I and 4O). These data suggest that endogenous Ct suppresses Hmx expression in C3da to a “basal” level, and overexpressed Ct in C2da mimics this effect and reduces Hmx to the basal level.

Reciprocally, we observed much higher Ct-GFP levels in ddaA (2.79-fold) and ddaF (2.85-fold) than in ddaB (Figures 4J and 4N), consistent with previous reports (*14*). *Hmx* KD strongly enhanced Ct-GFP in ddaB (136% increase) without affecting it in ddaA/F (Figures 4K and 4N). Finally, Hmx overexpression reduced Ct-GFP in ddaA/F to the basal level observed in ddaB (Figures 4L and 4P).

Together, these data suggest a stoichiometric competition model in which Ct and Hmx are “seeded” in newborn C2da and C3da neurons from their progenitors and mutually repress each other to establish an asymmetric dominance in each neuron. However, these dominances can be flipped by weakening the stronger factor or forcing high levels of the weaker one. Crucially, the “repressed” factor is not fully silenced but is maintained at a basal level. Thus, C2da and C3da identities are not defined by the absolute presence or absence of these factors, but by the stoichiometric asymmetric dominance established by active mutual repression. This high-tension equilibrium ensures that the latent, competing sensory program is continuously kept in check rather than permanently extinguished.

### The bistable switch remains malleable across development and into mature neurons

The above data show that the asymmetric dominance between Hmx and Ct establishes C2da/C3da identity. Is this imbalance subject to change only during a developmental critical period? Or do mature neurons require active Hmx/Ct repression to maintain their identity? To distinguish these possibilities, we used a temperature-sensitive Gal80 (*Gal80^ts^*) to restrict knockdown to defined temporal windows. Born at mid-embryogenesis and differentiating into functioning sensory neurons before hatching (22 h after egg laying, AEL), da neurons continue to expand dendric arbors during the larval period (∼98 h) (*29*). *ct* KD (at 29°C to inhibit Gal80^ts^) for only the first 24 h AEL was sufficient to switch marker expression in ddaA/F (Figures 5A and 5B). This conversion persisted through the 3^rd^ instar even after the KD was relieved (at 18°C) (Figure 5C). A reciprocal C2da-to-C3da conversion was observed for ddaB in both 1^st^ and 3^rd^ instar larvae when *Hmx* was similarly knocked down only in the first 24 h AEL (Figures 5D and 5E). These data suggest that once the switch is flipped during differentiation, the new state becomes self-sustaining.

**Figure 5.**
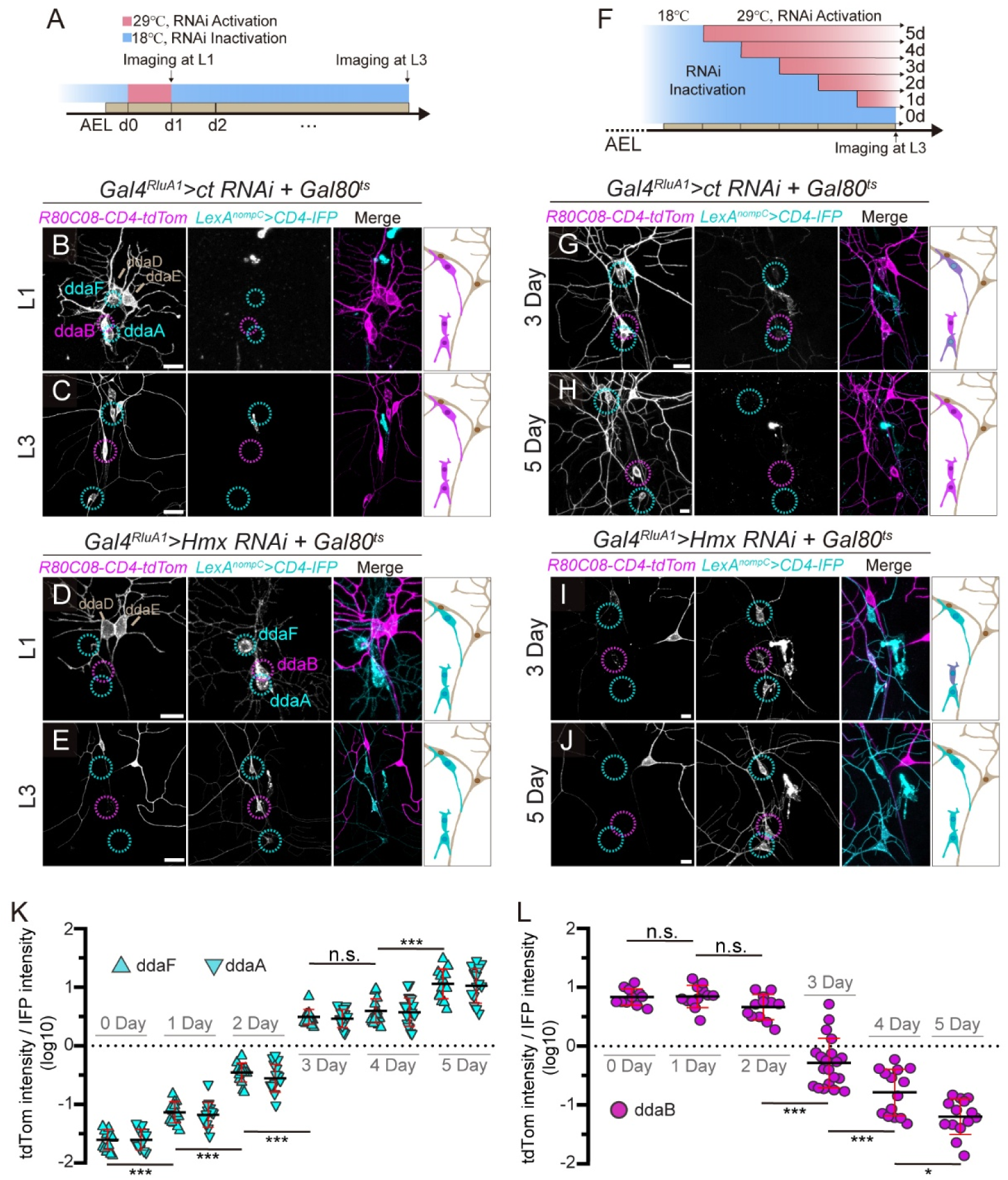
The bistable switch remains malleable across development and into mature neurons. **(A)** Experimental design for early and transient RNAi induction using the *Gal80^ts^* system. *Gal4^RluA1^* activity was restricted to the first 24 h after egg laying (AEL) (at 29°C) and suppressed for the rest of the larval period (at 18°C). **(B–E)** Expression of markers (*R80C08-CD4-tdTom* for C1da/C2da and *LexA^nompC^>CD4-IFP* for C3da) in da neurons following transient *ct* RNAi (**B** and **C**) or *Hmx* RNAi (**D** and **E**). Images were acquired at the L1 stage (immediately after induction) (**B** and **D**) and the late L3 stage (**C** and **E**). **(F)** Experimental design for variable durations of late RNAi induction with *Gal4^RluA1^* and *Gal80^ts^*. **(G–J)** Marker expression in da neurons following 3-day and 5-day induction of *ct RNAi* (**G, H**) or *Hmx RNAi* (**I, J**). RNAi expression is driven by *Gal4^RluA1^* under *Gal80^ts^* control (29°C for induction). In all image panels, cyan circles indicate C3da neurons (ddaA/F); magenta circles indicate C2da neurons (ddaB). Diagrams on the right illustrate cell conversion outcomes. Scale bars: 10 μm. **(K)** Ratio of marker intensities (*R80C08-CD4-tdTom*/*LexA^nompC^>CD4-IFP*) in ddaF (upward triangles) and ddaA (downward triangles) neurons with variable durations of *ct* KD. **(L)** Ratio of marker intensities in ddaB neurons with variable durations of *Hmx* KD. In (**K** and **L**), values are log_10_-transformed, and statistical comparisons are shown only for adjacent time points. In (**K**), ddaF and ddaA show no significant difference at any time point. Sample sizes (n = neurons): *ct RNAi* (n = 10-16 per timepoint); *Hmx RNAi* (n= 12-20 per timepoint). One-way ANOVA with Tukey post-hoc (*p<0.05, ***p<0.001, n.s., not significant). Black bars, mean; red bars, SD.

We then asked whether neurons still exhibit plasticity after initial differentiation. For this, we allowed neurons to differentiate normally (at 18°C) and initiated KD (at 29°C) at various larval stages (Figure 5F). We observed a cumulative conversion. Extending the duration of *ct* KD in larvae progressively increased the ratio of the C2da marker (tdTom) to the C3da marker (IFP) in ddaA/F (Figures 5G, 5H, 5K, S6A-S6D), with KD during strictly the final day of the larval development being sufficient to alter the ratio. *Hmx* KD showed a similar duration-dependent effect (Figures 5I, 5J, 5L, S6E-S6H), even though KD in the last 3 days was minimally required to change the markers. Together, these results establish that C2da/C3da identity is not a developmental endpoint but a continuously maintained state: even fully differentiated neurons remain poised to transdifferentiate into the competing identity once repression of the alternative selector is lost.

### Neuronal reprogramming extends to axon projections and sensory function

Having established that the Ct/Hmx switch is continuously required to maintain class-specific transcriptional programs in mature neurons, we next asked whether altering this switch can affect neuronal features specified at much earlier developmental stages. Da neurons establish class-specific axon projection patterns in the central nervous system (CNS) by embryonic stages 16-17 (*30*), well before the establishment of dendritic arbors. We thus examined the effects of *ct* and *Hmx* manipulations on axon projection and organismal sensory behavior. To achieve the stringent, single-cell resolution required to trace overlapping axonal projections in the dense ventral nerve cord, we utilized MAGIC (Mosaic Analysis by gRNA-Induced Crossing-over) (*31–33*) to generate single neuronal clones expressing *Gal4^21-7^>CD4-tdTom*. Clone identities were determined based on soma locations, dendrite morphologies, and overlap or exclusion with *R80C08-CD4-tdGFP* (Figure 6A). Due to the lineage-specific recombination frequencies inherent to the MAGIC system, we utilized ddaA as the representative C3da neuron for axonal analysis. Wildtype (WT) ddaB (C2da) axons consistently terminated in an inverted “T” shape, with variable lengths of the “T” arms (Figures 6B, S7A-7D). In contrast, WT ddaA (C3da) axons exhibited an “L” shape (Figures 6C, S7E-S7H). Strikingly, *Hmx* KD in ddaB clones transformed the axonal trajectory from the inverted “T” to the “L” shape in 3 out of 5 neurons (Figures 6D, S7I-S7L). Given the short temporal window within which Gal4/UAS-driven KD must take effect before axons reach the CNS, these data strongly suggest that the Ct/Hmx switch controls axon projection as well.

**Figure 6.**
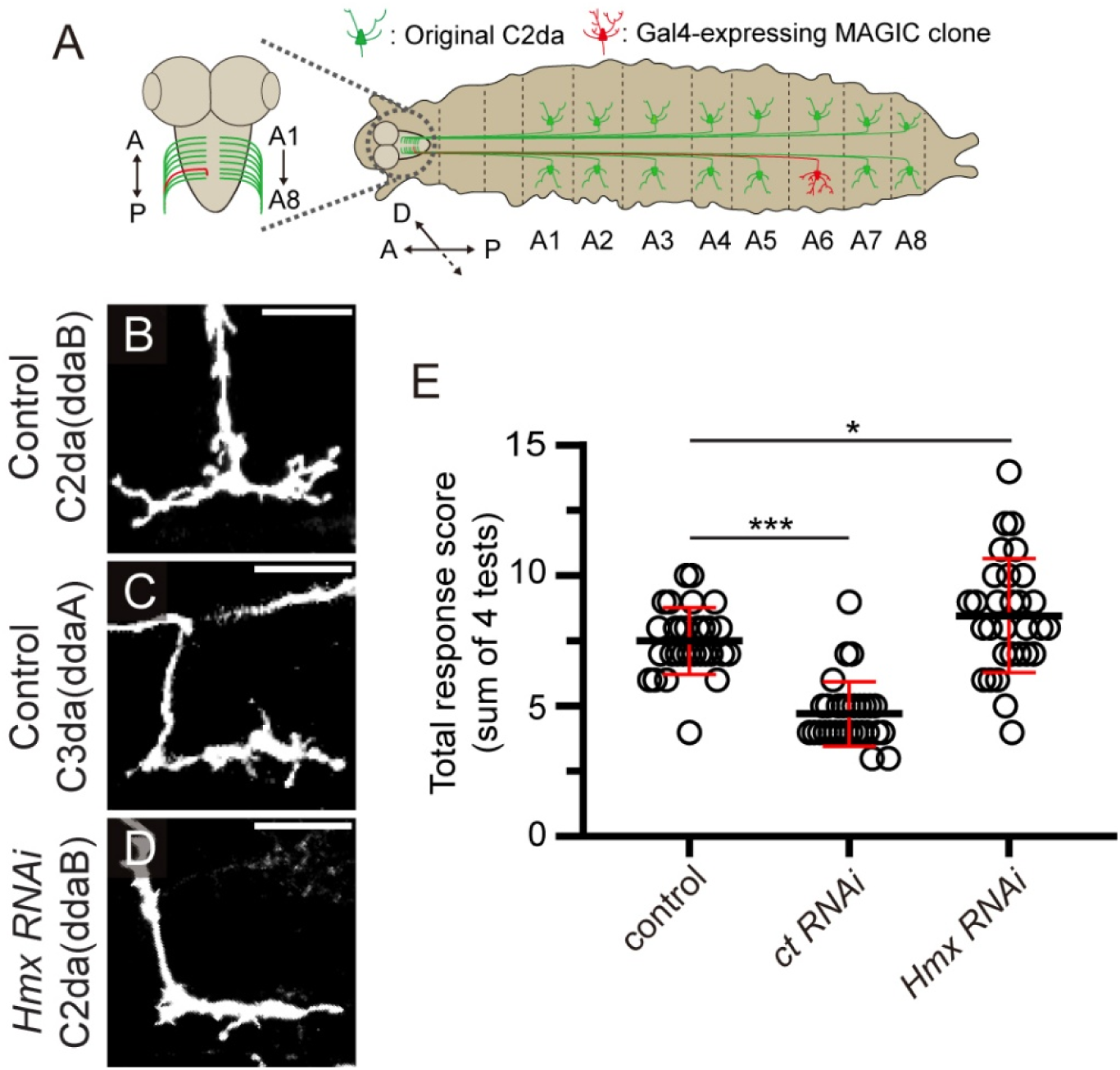
Neuronal reprogramming extends to axon projections and sensory function. **(A)** Diagram of gene knockdown in sparse neuronal clones generated by the MAGIC technique, with C2da neurons as an example. C2da neurons are labeled by a green marker, while MAGIC clones express a red marker. The identity of a clone is determined by the soma position and dendrite morphology, facilitated by the C2da marker. After a proper clone was identified, its axon projection in the VNC was examined. **(B–P)** Axon terminal morphologies of WT ddaB (C2da) (**B–F**) and ddaA (C3da) (**G–K**) clones and *Hmx*-KD ddaB clones (**L–P**). The MAGIC clones were induced by *zk-nCas9(H840A)* and *gRNA-81F(Gal80)*; the clones were labeled by *Gal4^21-7^>UAS-CD4-tdTom*. Scale bars: 15 μm. (**Q**) Diagram of the gentle touch assay and the scoring scheme (0–4), adapted from (*12*). (**R**) Total response scores (sum of 4 tests) across indicated genotypes. RNAi was expressed by *Gal4^RluA1^*. Sample sizes (N = animals): Control (N = 30); *ct RNAi* (N = 30); *Hmx RNAi* (N = 30). One-way ANOVA with Tukey post-hoc (*p<0.05, ***p<0.001). Black bars, mean; red bars, SD.

Finally, to test whether cell fate conversion has organismal consequences, we tested the larval response to gentle touch, a behavior mediated specifically by C3da neurons (Figure S7M) (*12*). *ct* KD, which converts touch-sensitive C3da neurons into C2da-like neurons, significantly reduced the behavioral response (Figure 6E). Conversely, *Hmx* KD, which generates supernumerary C3da-like neurons, resulted in hypersensitivity to gentle touch (Figure 6E). Thus, manipulating the C2da/C3da toggle switch does not merely alter cellular morphology, but functionally reprograms axon projections and alters organismal behavior, suggesting that this logic gate dictates sensory modality.

### Human homologs of Ct and Hmx possess conserved activities that toggle sensory identity

Ct and Hmx belong to ancient CUX and HMX homeodomain families. By analyzing human dorsal root ganglion (DRG) transcriptomic data (*34*), we observed overlapping expression patterns of *CUX1* and *HMX1* in multiple sensory neuron types and *CUX2* expression in a subset of these neurons (Figure S8A). Using 313 orthologous genes comprising ion channels, transcription factors, signaling molecules, and cell adhesion proteins (Supplementary Table 2) to conduct MetaNeighbor analysis (*35*) between *Drosophila* da neurons and human DGR neurons (Figure 7A), we found that C2da neurons exhibit a molecular signature resembling human Aδ High-Threshold Mechanoreceptors (Ad HTMRs). Conversely, C3da neurons share the highest degree of transcriptomic similarity with human cold-sensing nociceptors and Aδ Low-Threshold Mechanoreceptors (Ad LTMRs), which detect hair movement and gentle touch (*36*). These similarities are particularly obvious in the expression of subtype-specific channels, cytoskeletal and signaling regulators (Figure 7B). Notably, *CUX2* is expressed in an Aδ sublineage of sensory neurons in mice, and its LOF causes mechanical hypersensitivity (*37*), consistent with an expansion of Aδ HTMRs at the expense of Aδ LTMRs. The transcriptomic resemblances between *Drosophila* and human sensory neurons suggest that the functional gene modules defining these fly and human sensory subtypes may share a conserved genomic blueprint.

**Figure 7.**
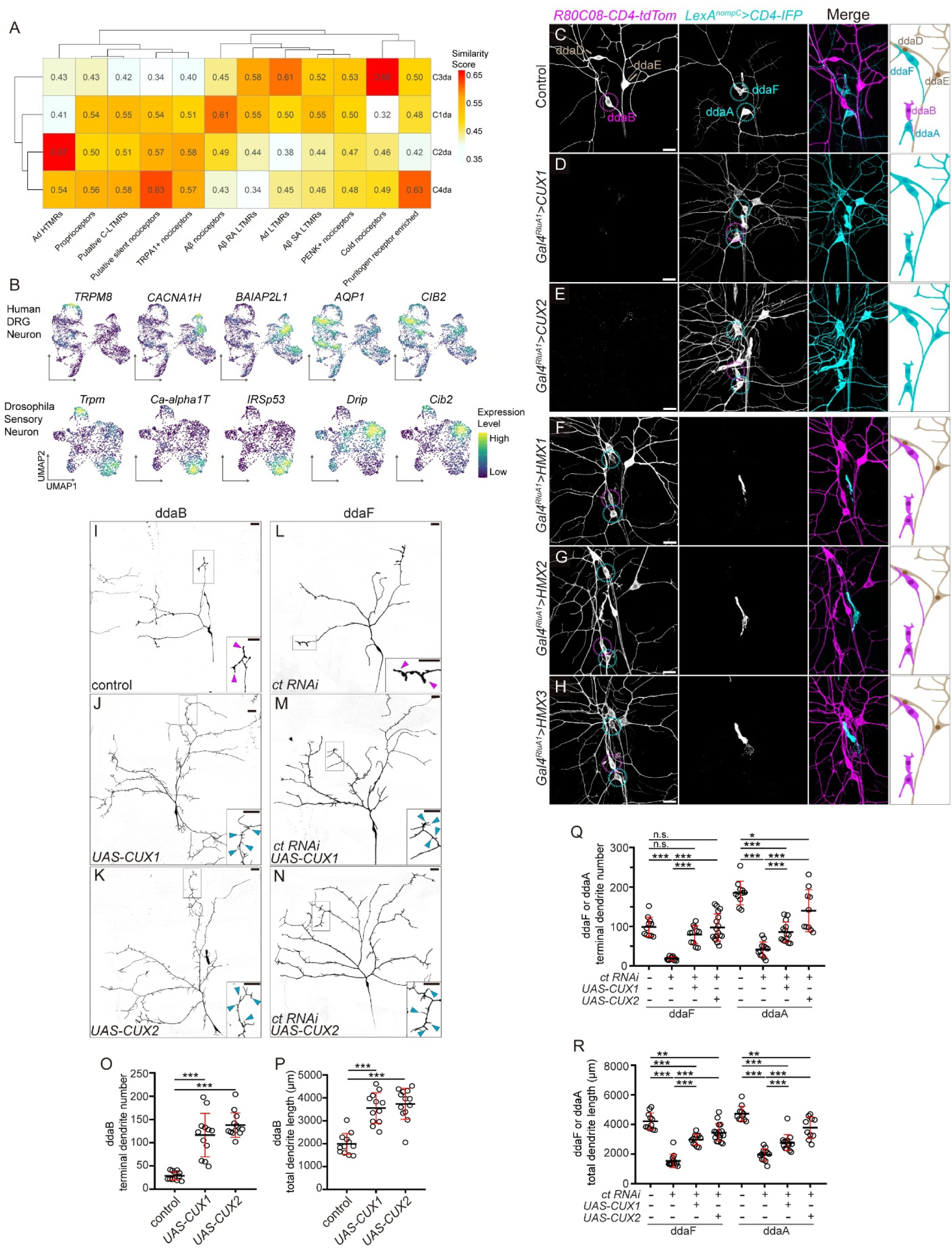
Human homologs of Ct and Hmx possess conserved activities that toggle sensory identity. **(A**) MetaNeighbor analysis between major clusters of human DRG neurons and *Drosophila* da neurons based on expression of 313 pairs of orthologous genes that are differentially expressed in DRG and da neurons. The color indicates the degree of similarity. **(B)** UMAPs of human DRG and *Drosophila* da neurons, showing expression of select pairs of orthologous genes. TRPM8 and Trpm are calcium channels in the TRPM family involved in cold sensing. CACNA1H and Ca-alpha1T are low-voltage–activated calcium channels. BAIAP2L1 and IRSp53 are BAR/IMD-domain–containing adaptor protein involved in actin cytoskeleton remodeling. AQP1 and Drip are water channel proteins. CIB2 and Cib2 are calcium- and integrin-binding protein containing EF-hand domains. **(C–H)** Expression of markers (*R80C08-CD4-tdTom* for C1da/C2da and *LexA^nompC^>CD4-IFP* for C3da) in late 3rd instar larvae of WT control (**C**), *CUX1* OE (**D**), *CUX2* OE (**E**), *HMX1* OE (**F**), *HMX2* OE (**G**), and *HMX3* OE (**H**). OE was driven by *Gal4^RluA1^* and restricted to the last 72 h (for *CUX1* OE and *HMX1* OE) or 96 h (for *HMX2* OE and *HMX3* OE) using *Gal80^ts^* to bypass early larval lethality. Cyan circles indicate C3da (ddaA/F); magenta circles indicate C2da (ddaB). The diagrams on the right illustrate cell conversion outcomes. Scale bars: 10 μm. **(I-N)** CoinFlp clones of ddaB (C2da) neurons in WT control **(I)** or expressing *UAS-CUX1* **(J)** or *UAS-CUX2* **(K)**. **(L-N)** CoinFlp clones of ddaF neurons expressing *ct* RNAi alone **(L)**, or co-expressing *UAS-CUX1* **(M)** or *UAS-CUX2* **(N)**. In **(I-N)**, insets show high-magnification views of dendritic terminals. Cyan arrowheads indicate dendritic spikes; magenta arrowheads indicate varicosities. Scale bars: 25 μm. **(O and P)** Terminal dendrite number (**O**) and total dendrite length (**P**) of ddaB neurons. Sample sizes (n = neurons, N = animals): Control (n = 12, N = 12); *UAS-CUX1* (n = 12, N = 12); *UAS-CUX2* (n = 13, N = 13). **(Q and R)** Terminal dendrite number (**Q**) and total dendrite length (**R**) in indicated genotypes. Sample sizes: ddaF_*ct RNAi* (n = 12, N = 12); ddaF_*ct RNAi* + *CUX1* (n = 12, N = 12); ddaF_*ct RNAi* + *CUX2* (n = 17, N = 15). One-way ANOVA with Tukey post-hoc (*p<0.05, **p<0.01, ***p<0.001; n.s., not significant). Black bars, mean; red bars, SD.

To test functional conservation of the CUX and HMX families, we expressed human *CUX1/2* and *HMX1/2/3* in fly neurons. Overexpression of *CUX1* and *HMX1/2/3* in all da neurons resulted in early larval lethality, but attenuating the expression with Gal80^ts^ extended the viability to late larval stages. Human *CUX* genes mimicked fly *ct*: their expression caused marker switching in C1da (ddaD/E) and C2da (ddaB) neurons (Figures 7C-7E). Conversely, overexpression of each *HMX* gene induced opposite marker switching in C3da (ddaA/F) neurons (Figures 7F-7H), mirroring *Hmx* overexpression.

Next, we asked whether the human genes can also transform dendrite arbors of individually labeled da neurons like their *Drosophila* homologs. Indeed, coinFlp clones of ddaB expressing either *CUX1* or *CUX2* resembled WT ddaF morphology, reflected by increased dendrite length and branch number, gain of dendritic spikes, and simultaneous loss of local varicosities (Figures 7I-7K, 7O, 7P). CoinFlp clones expressing any of the *HMX* genes showed severe dendrite degeneration, likely due to high potencies of these TFs, precluding dendrite quantification. Finally, we asked whether *CUX* genes can functionally replace *ct* in C3da neurons. While expressing *ct-RNAi* in coinFlp clones of ddaF and ddaA caused the neurons to adopt C2da arbors (Figures 7L and S8B), co-expressing *CUX1* or *CUX2* with *ct-RNAi* in these neurons restored the C3da dendrite morphologies (Figures 7L-7N, 7Q, 7R, S8C, S8D).

Together, the above results demonstrate that human CUX and HMX genes possess similar biological activities as their *Drosophila* homologs, capable of interconverting *Drosophila* C2da and C3da neurons. Thus, they retain the ability to execute this mutually exclusive logic gate and function as opposing terminal selectors in *Drosophila* sensory neurons.

## DISCUSSION

### A bistable switch of mutually repressive terminal selectors maintains discrete neuronal identities

Our findings establish that the terminal identities of *Drosophila* C2da and C3da somatosensory neurons are not static developmental endpoints, but dynamic states sustained by the continuous competition of two opposing terminal selectors. Ct acts as a dual-function “C3da Selector/C2da Repressor,” while Hmx acts as a “C2da Selector/C3da Repressor” (Figure 8A). These two transcription factors each execute a tripartite logic: they simultaneously activate their specific effector genes, sustain their own expression through positive feedback, and repress the opposing selector (Figure 8A). This generates a robust bistable switch with two discrete, self-stabilizing states. Loss of either factor in fully differentiated neurons triggers rapid, deterministic acquisition of the alternative fate, demonstrating that identity maintenance is an active process, not a passive consequence of stable chromatin or transcriptional programs.

**Figure 8.**
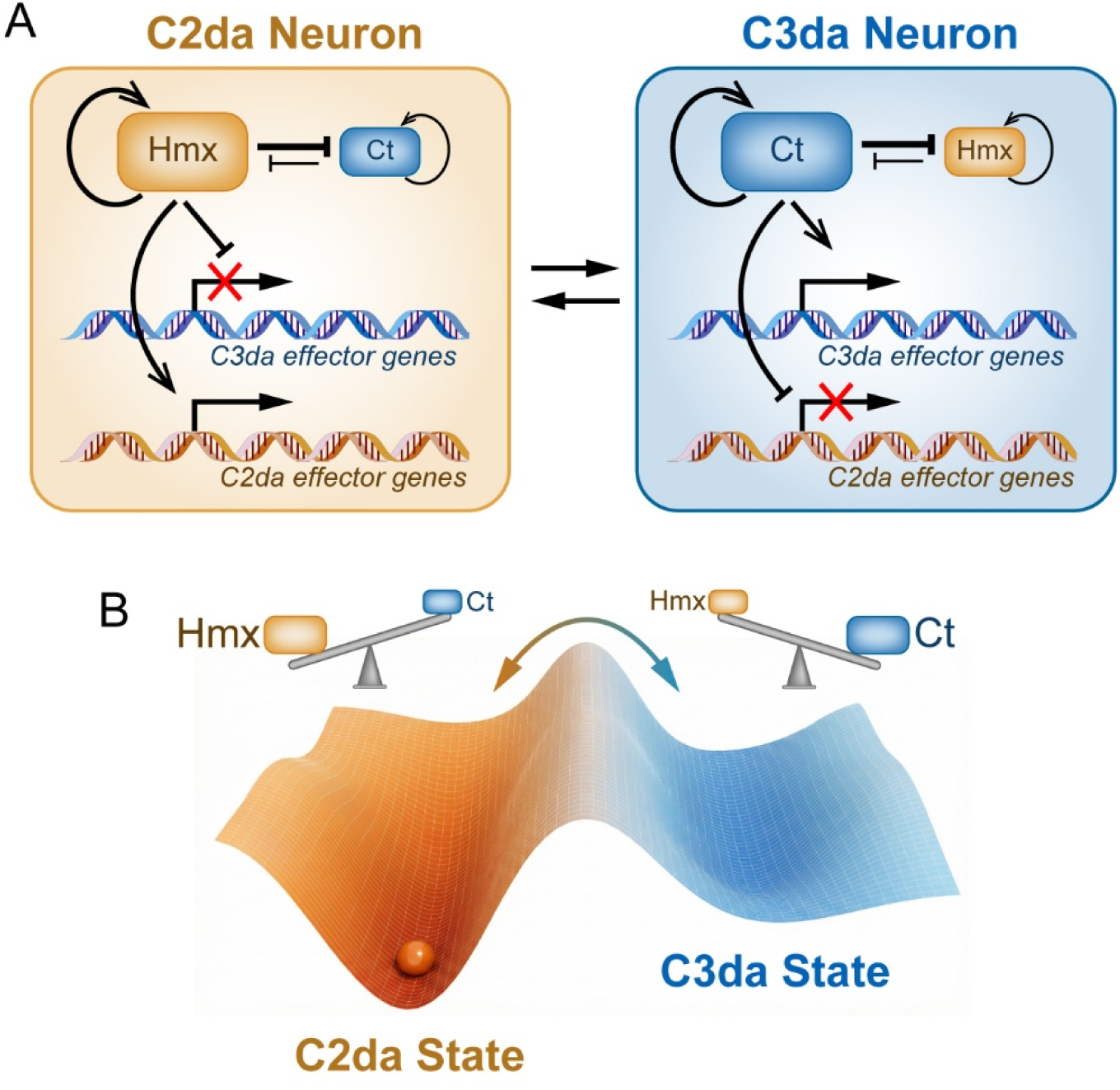
Cut and Hmx form a bistable switch to safeguard neuronal identity via a “see-saw” mechanism. **(A**) Diagram illustrating the tripartite logic of Hmx and Ct as opposing terminal selectors in C2da and C3da neurons. The model depicts how this bistable switch operates via the simultaneous regulation of class-specific effector genes, positive autoregulation to amplify and lock the dominant state, and mutual cross-repression to suppress the alternative identity. **(B)** A “see-saw” model for Hmx and Ct to maintain mature states of C2da and C3da neurons in a high-tension energy landscape. C2da and C3da identities are two metastable states established by the asymmetric dominance between Hmx and Ct, and C2da identity is the default state in the absence of terminal selectors. The imbalance between these two factors can be flipped to establish the opposite dominance, resulting in identity interconversion.

This mechanism differs in an important way from the prevailing view of terminal selectors. In most established systems, terminal selectors lock in cell fate primarily through positive regulation of effector batteries and autoregulatory feedback, with repression of alternative programs seen as a secondary safeguard against transcriptional noise (*2, 6*). The Ct/Hmx circuit operates by a fundamentally different logic: the terminal selectors themselves antagonize each other in a symmetric, mutually repressive loop. This cross-repression is not a peripheral feature but the primary instructive force defining cellular identity. Interestingly, in the absence of both factors, post-mitotic neurons default to a C2da-like ground state (Figure 8B), revealing that Hmx’s essential instructive function is to repress Ct and thereby prevent the C2da program from being overwritten by the Ct-driven C3da program. Thus, identity in this system is defined not by selector presence, but by selector dominance.

Cross-repressive transcription factors are known to partition spatial domains of neural progenitors in the developing nervous system, exemplified by the homeodomain code that subdivides the vertebrate ventral neural tube (*38*). Our results demonstrate that this regulatory design is not restricted to transient developmental decisions in proliferating progenitors, but extends post-mitotically to defend the identities of mature neurons throughout life. In doing so, the Ct/Hmx switch resolves a long-standing question about terminal selector function: whether cross-repression is merely a noise-dampening accessory or a primary, instructive force. Our data argue decisively for the latter.

### Asymmetric dominance seeded by developmental history sustains the bistable state

Our protein-level data reveal that bistability is achieved by asymmetric dominance: each mature neuron expresses both Ct and Hmx, with the dominant factor present at substantially higher levels. The repressed factor is not silenced, but rather held at a basal level, and forced overexpression of the subordinate factor is sufficient to flip the dominance and switch the neuronal identity. We refer to this configuration as a “see-saw” (Figure 8B), in which identity is determined by which factor dominates rather than which factor is present.

This arrangement places mature neurons in a state of high-tension metastability: the latent program is never extinguished but continuously suppressed by its competitor. However, when the dominant factor is weakened, the latent factor is rapidly de-repressed and amplified through autoregulation, tipping the see-saw and committing the cell to the alternative fate. The system is thus simultaneously optimized for fidelity (prevention of stochastic switching by the steep energy barrier) and responsiveness (tipping the balance by a single perturbation).

How is the correct initial state of the switch chosen? Our pre-mitotic LOF analysis reveals an asymmetric co-requirement for Ct and Hmx in both lineages, explaining their dual inheritance in post-mitotic neurons. We propose a “seed and lock” model in which the distinct developmental histories of adjacent proneural clusters establish an initial stoichiometric imbalance in newborn neurons, biasing the see-saw toward one fate, and the mutual repression loop then translates this transient, lineage-derived asymmetry into a permanent, life-long identity. Thus, lineage history acts not as a deterministic instructor of fate but as a bias-setter, with post-mitotic mutual repression providing the lock.

Stoichiometric competition between transcription factors is a recurring principle in cell fate decisions, most famously between PU.1 and GATA1 during hematopoietic commitment (*39*). The Ct/Hmx switch extends this principle into the post-mitotic domain, showing that stoichiometric competition can govern not only the initial fate decision but its lifelong defense.

### A potentially conserved logic gate for segregating somatosensory modalities

Three lines of evidence suggest that this regulatory module is evolutionarily conserved. First, cross-species transcriptomic comparison aligns C2da neurons with human Aδ HTMRs, and C3da neurons with both Aδ LTMRs and cold-sensing nociceptors. The latter dual mammalian alignment mirrors the known multimodal function of C3da neurons (*12, 13*), suggesting that ancestral sensory modalities co-deployed in insect neurons were subsequently partitioned across specialized mammalian subtypes. Second, *CUX1* and *HMX1* are co-expressed in multiple human DRG neuron subtypes (Figure S7A), satisfying the prerequisite for a mutually repressive switch. Third, and most directly, human CUX1/2 and HMX1/2/3 functionally substitute for their *Drosophila* counterparts, with CUX1/2 also rescuing the dendritic and identity defects of *ct* loss. Thus, despite >500 million years of divergent evolution, these proteins retain the biochemical activities required to participate in this switch.

Complementary mouse genetics support this possibility: *Cux2* loss in an Aδ sublineage causes mechanical hypersensitivity (*37*), consistent with an HTMR-biased shift predicted by the see-saw model. Although direct mutual repression between mammalian CUX and HMX paralogs remains to be tested, the convergence of expression, function, and rescue data suggests that this regulatory logic predates the divergence of protostomes and deuterostomes. We propose that the CUX/HMX module represents an ancient genomic blueprint for partitioning somatosensory modalities, which was subsequently elaborated in vertebrates through gene duplication and combinatorial deployment with other selectors. If so, in mammalian DRG neurons co-expressing CUX and HMX paralogs, perturbation of one paralog should drive transcriptomic and functional drift toward the alternative subtype, a prediction made readily testable by existing mutant alleles.

### Implications for cellular identity, plasticity, and disease

The capacity of mature C2da and C3da neurons to undergo deterministic, lineage-crossing transdifferentiation challenges the view of terminal differentiation as an irreversible endpoint. Where terminal selector loss in other systems typically produces hybrid states or partial identity erosion (*2, 5*), the Ct/Hmx circuit operates as a discrete, high-fidelity toggle that replaces the whole identity, across transcriptome, dendrites, axons, and behavior. The epigenetic barriers separating sensory lineages are therefore not deeply entrenched chromatin states but labile equilibria continuously enforced by the stoichiometric balance of competing selectors.

While pioneering work achieved direct lineage reprogramming via forced transcription factor expression (*40–42*), our *in vivo* results show that complete interconversion is a latent capability of mature neurons, gated by a single endogenous switch. Critically, our results indicate this transdifferentiation occurs directly, without reversion to a dedifferentiated or progenitor-like state. This raises the possibility that other apparently terminal cell types harbor similar latent switches that could be exploited for targeted *in vivo* manipulation without passage through a pluripotent intermediate.

The same metastability also exposes a vulnerability in disease. *CUX1* and *HMX1* mutations are linked to neurodevelopmental and sensory disorders (*43, 44*), typically interpreted as defects in specification or survival. Our model suggests an additional possibility: partial loss-of-function may shift the see-saw without abolishing it, producing neurons of intermediate identity, an “identity drift” that could underlie sensory processing abnormalities. In addition, age-related decline in maintenance factor expression could compromise subtype identity in neurons that have functioned correctly for decades.

Together, our findings address a fundamental question in cellular and systems biology: how do complex tissues maintain functional fidelity when constructed from cells with overlapping transcriptional histories? The Ct/Hmx switch provides one answer: discrete identity is not a static developmental endpoint, but a poised, reversible equilibrium actively defended through continuous competition between opposing fate determinants. This active, competitive logic may represent a general principle by which biological systems segregate and maintain discrete cellular states throughout the lifespan of the organism.

## MATERIALS AND METHODS

### Drosophila strains

The details of fly strains and reagents used in this study are listed in the Key Resource Table (Supplementary Table 3). Flies were grown on standard yeast-glucose medium at 25°C under a 12:12 h light/dark cycle, unless otherwise specified. For post-mitotic gene manipulation in da neurons, we used pan-da drivers *Gal4^RluA1^* (*20*) and *Gal4^21-7^* (*19*). Unless noted otherwise, neuronal subtypes were distinguished using *R80C08-CD4-tdTomato* (labeling C1da and C2da), *LexA^NompC^ > CD4-IFP* (labeling C3da) (*17*), and *ppk-CD4-tdGFP* (labeling C4da) (*16*). Endogenous protein expression was examined using *Ct-GFP* and *Hmx-GFP* knock-in lines. *SOP-Cas9* (*27*) was used for early pre-mitotic KO in the neuronal lineages; *ase-Cas9* was used for KO later in the lineages and to reveal LOF phenotype in post-mitotic neurons.

See Supplemental Methods for detailed protocols on molecular cloning and generation of transgenic flies, generation of Trojan exon Gal4 reporters, live imaging, image analysis and morphological quantification, MAGIC-based sparse axonal labeling, gentle touch behavioral assays, snRNA-seq and data analysis, cross-species transcriptomic comparisons, and statistical analysis.

## ACKNOWLEDGMENTS

We thank Haiyuan Yu, Addgene, and the *Drosophila* Genomics Resource Center (DGRC) for plasmids; Yuh-Nung Jan, Marc Freeman, Ben White, the Bloomington *Drosophila* Stock Center, the KYOTO *Drosophila* Stock Center for fly stocks; Elizabeth Korn, Payton Ditchfield, and Christina Breneman for larval dissection; Colleen McLaughlin, Boxuan Zhao, Jen Grenier, and Kuang-Tse Wang for strategy and troubleshooting of snRNA-seq; Jen Grenier, Peter Schweitzer, Cornell Flow Cytometry Facility, and Cornell Genomics Facility for facilitating snRNA-seq; Shihong Gao, Hongjie Li, and Ayshwarya Subramanian for advice on analysis of snRNA-seq data; Patrick Murphy, Claude Desplan, and Oliver Hobert for feedback. This work was supported by NIH grants (R01NS099125, R24OD031953, and R01NS138421) awarded to C.H..

This manuscript is the result of funding in whole or in part by the National Institutes of Health (NIH). It is subject to the NIH Public Access Policy. Through acceptance of this federal funding, NIH has been given a right to make this manuscript publicly available in PubMed Central upon the Official Date of Publication, as defined by NIH.

## FUNDING

R01NS099125 (C.H.)

R24OD031953 (C.H.)

R01NS138421 (C.H.)

## AUTHOR CONTRIBUTIONS

**Conceptualization:** Qi-Nan Chen, Yineng Xu, Chun Han

**Data curation:** Qi-Nan Chen, Yineng Xu

**Formal analysis:** Qi-Nan Chen, Yineng Xu

**Funding acquisition:** Chun Han

**Investigation:** Qi-Nan Chen, Yineng Xu, Yifan Shen, Nambita S. Sahai

**Methodology:** Qi-Nan Chen, Yineng Xu, Chun Han

**Supervision:** Qi-Nan Chen, Yineng Xu, Chun Han

**Resources:** Qi-Nan Chen, Bei Wang

**Validation:** Qi-Nan Chen

**Visualization:** Qi-Nan Chen, Yineng Xu

**Writing – original draft:** Chun Han, Qi-Nan Chen

**Writing – review & editing:** Chun Han, Qi-Nan Chen, Yineng Xu

## COMPETING INTERESTS

The authors declare no competing interests.

## DATA AND MATERIALS AVAILABILITY

snRNA-seq data are available from the Gene Expression Omnibus (GEO) repository (accession number GSE324368). All other data generated or analyzed during this study are included in the manuscript and supporting files.

## SUPPLEMENTARY METHODS

### Molecular cloning and transgenic flies

#### R80C08-CD4-tdTom and R80C08-CD4-tdGFP

The R80C08 enhancer was PCR-amplified from *R80C08-Gal4* genomic DNA using oligos ggggACAAGTTTGTACAAAAAAGCAGGCTatagggtgagctg and ggggACCACTTTGTACAAGAAAGCTGGGTGGAATAGGG. The resulting DNA fragment was combined with pDONR-221 (Thermo Fisher Scientific) to generate pENTR221-R80C08 through a Gateway BP reaction (Thermo Fisher Scientific). pENTR221-R80C08 was then combined with destination vectors pDEST-HemmarR2 (Addgene 112813) (*27*) and pDEST-HemmarG2 (Addgene 112814) (*27*) to generate the expression vectors pHemmarR2-R80C08 and pHemmarG2-R80C08, respectively, through Gateway LR reactions (Thermo Fisher Scientific).

#### LexAOP2-CD4-IFP2.0-2A-HO1

The XhoI-PacI fragment of pDEST-APLO (Addgene 112805) (*45*) was replaced by the XhoI-PacI fragment of pIHEU-CD4-IFP2.0-HO1 (*46*) to make pAPLO-CD4-IFP2.0-HO1.

#### ss-CD4-tdTom

The *ss* enhancer was PCR-amplified from *w^1118^*genomic DNA using oligos ggggACAAGTTTGTACAAAAAAGCAGGCTAAAGTGACTAGGGGCTAAAAAG and ggggACCACTTTGTACAAGAAAGCTGGGTAATAAAGCCCACCCCCAG. The resulting DNA fragment was combined with pDONR-221 to generate pENTR221-ss through a Gateway BP reaction. pENTR221-ss was then combined with the destination vector pDEST-HemmarR2 to generate the expression vector pHemmarR2-ss through a Gateway LR reaction.

#### ase-Cas9

The *ase* enhancer was PCR-amplified from *w^1118^*genomic DNA using oligos ttgtgctcggcaacagcatGCTAGCGTCAATTCCGTTTACTCATC and CTTGTCCGAGGTACCCATAATTAAGTTTTTTGATTCGTG. The Cas9 coding sequence (CDS) was PCR-amplified from pHACK(Gal4)-DONR(T2A-Cas9) (Addgene 194768) (*47*) using oligos CTTAATTATGGGTACCTCGGACAAGAAGTACTCCATTGG and CCGACTGGCTTAGTTAattaatttctagaTTAGGCGTAGTCTGGGACG. Both fragments were assembled into SphI/XbaI digested pAPIC-nsyb-tdGFP (*27*) using NEBuilder HiFi DNA assembly (New England Biolabs) to make pAPIC-ase-Cas9.

#### UAS-Hmx-IFP

The Hmx coding sequence (CDS) was PCR amplified from DGRC clone FMO10659 using oligos actctgaataGGGAATTGGGAATTcagaaaATGTCTTCCTCCGAGGCCGA and CGTCGTCGTCTTTGTAGTCGCTAGCGCTTGCTACCAGCGACGAG. The resulting fragment was assembled into EcoRI/NheI-digested pACU-Eato(C)-FLAG (*48*) by NEBuilder HiFi DNA assembly to make pACU-Hmx-FLAG. The emiRFP670 CDS was PCR-amplified from pLAMP1-emiRFP670 (Addgene 136570) with oligos cttcGCTAGCttcggctccaccggctc and ctctCTCGAGGCTCTCAAGCGCGGTG and then digested by NheI and XhoI. The T2A-HO1 CDS was PCR amplified from pDEST-HemmarI2 (Addgene 112815) (*27*) using oligos AGGGTCTGGTGGCAGCGGActcGAGGGCAGAGGAAGTC and CCGACTGGCTTAGTTAattaattctagaaTCACATAGCATAGAGCCCC and then digested by XhoI and XbaI. Both emiRFP670 and T2A-HO1 fragments were then ligated into NheI/XbaI-digested pACU-Hmx-FLAG to make pACU-Hmx-emiRFP670-T2A-HO1.

#### Ct-GFP KI construct

Two *ct* gRNA target sequences (TTTACTTTAGTAGTTCCAAC and CAGCAACTGCGGCAGCCGGT) were incorporated into PCR primers and cloned into pAC-CR7T-gRNA2.1-nlsBFP (Addgene 170515) by following published protocols (*47*) to make pACGCB(2.1)-cut. Four fragments amplified by PCR were assembled into PstI/NheI digested pACGCB(2.1)-cut by NEBuilder HiFi DNA assembly to generate the final KI vector pHACK(cutC)-DONR(sfGFP-2A-QF2). The four fragments are superfolder GFP (sfGFP) CDS amplified from pIHEU-sfGFP-LactC1C2 (*49*) using oligos CGGCAGCCGGaTGGAACTACGGATCCGGTGGCGGCGGAAG and CCTCTGCCCTCGGAAGATCTCTTATACAGCTCGTCCATGCCCAG, T2A-QF2 CDS amplified from *ppk-QF2* genomic DNA using oligos AGATCTTCCGAGGGCAGAGGAAGCCTGCTGACCTGCGGCGATG and tttttttactTcACTtcTtGTAcGTgTTAATGTCGGAGAAGTTACATCCAGGAACGAATC, *ct* 5’ homology arm (HA) amplified from *w^1118^* genomic DNA using oligos gtacgcaAAGCTTGGCTGCATGAGCGGTGCCCCCATTCCCTATC and GTAGTTCCAtCCGGCTGCCGCAGTTGCTG, and *ct* 3’ HA amplified from *w^1118^*genomic DNA using oligos cACgTACaAgaAGTgAagtaaaaaaattaacagaactaaatataaattcagtgcaatgtgtagag and TTAGCGACGTGTTCACTTTGTTTATTTGGGCAACAAAATAGGGGAAAACTTTTCTGC.

#### Hmx-GFP KI vector

Two *Hmx* gRNA target sequences (CGTCGCTGGTATAGGGAGTG and CCCTATACCAGCGACGAGAG) were incorporated into PCR primers and cloned into pAC-CR7T-gRNA2.1-nlsBFP to make pACGCB(2.1)-Hmx. Three fragments amplified by PCR were assembled into PstI/NheI digested pACGCB(2.1)-Hmx by NEBuilder HiFi DNA assembly to generate the final KI vector pHACK(HmxC)-DONR(sfGFP-2A-QF2). The three fragments are a sfGFP-T2A-QF2 fragment amplified from pHACK(cutC)-DONR(sfGFP-2A-QF2), *Hmx* 5’ homology arm (HA) amplified from *w^1118^* genomic DNA using oligos gtacgcaAAGCTTGGCTGCAATAATAACAACAGCAACACCACTG and CTTCCGCCGCCACCGGATCCTACCAGgGAgctGAGCGGGGGTCTCGGGGGGCTGGTGT TG, and *Hmx* 3’ HA amplified from *w^1118^* genomic DNA using oligos CATTAAcACgTACaAgaAGTgAGGAGTGGGGTGCCTGGCGAC and TTAGCGACGTGTTCACTTTGtatcgaacttagcctacacttgac.

#### gRNA-ct, gRNA-Hmx, and gRNA-Ct-Hmx

*ct* gRNA target sequences (TATCACTACCAAGATCAAGG and ATGAGCTGGGCTTGGCAACG) and *Hmx* gRNA target sequences (GGAGGGCGAGGAGATCATCG and GTAGCGCTTCAGATCGAACG) were cloned in pairs (for *ct* and *Hmx* single gene KO) or all together (for *ct*/*Hmx* double KO) into pAC-U63-gRNA2.1 (Addgene 170512) by following published protocols (*47*).

#### UAS-hCUX1-FLAG

The human *CUX1* CDS was PCR-amplified from Horizon MGC cDNA (clone ID: 5740343) with oligos actctGAATAGGGAATTGGGAATTCaaaATGGCGGCCAATGTGGGATC and CGTCGTCGTCTTTGTAGTCgctagcGAACTCCCATTCGATAGGTTCCTC. The fragment was cloned into EcoRI/ NheI-digested pACU (Addgene 58372) (*16*) by NEBuilder HiFi DNA assembly to make pACU-hCUX1-FLAG.

#### UAS-hCUX2-FLAG

The human *CUX2* CDS is PCR-amplified from Horizon MGC cDNA (clone ID: 100014492) with oligos actctGAATAGGGAATTGGGAATTCaaaATGGCCGCCAATGTGGG and CGTCGTCGTCTTTGTAGTCgctagcGAACTCCCACTCCAGGGC. The fragment was cloned into EcoRI/ NheI digested pACU by NEBuilder HiFi DNA assembly to make pACU-hCUX2-FLAG.

#### UAS-hHMX2-FLAG

The human *HMX2* CDS is PCR-amplified from a custom cDNA clone (a gift from Haiyuan Yu) with oligos actctGAATAGGGAATTGGGAATTCaaaATGGGCAGCAAAGAAGATGC and CGTCGTCGTCTTTGTAGTCgctagcGTAGTCGAGCTTGTTGTATAGGTTG. The fragment was cloned into EcoRI/ NheI digested pACU by NEBuilder HiFi DNA assembly to make pACU-hHMX2-FLAG.

#### UAS-hHMX1-FLAG and UAS-hHMX3-FLAG

The hHMX1 and hHMX2 CDS were synthesized by GeneScript and cloned into the pACU vector.

Transgenic constructs were injected by GenetiVision and Rainbow Transgenic Flies to transform flies through φC31 integrase-mediated integration into attP docker sites.

### Generation of *ct-GFP* and *Hmx-GFP* KI strains

To make *ct-GFP*, pHACK(cutC)-DONR(sfGFP-2A-QF2) was injected into embryos of *y w; nos-Cas9(y+)* by GenetiVision. The injected animals were crossed to *QUAS-6XGFP*. GFP-positive animals were selected from their progeny at larval stages and later crossed to the *FM6* balancer to separate the *ct-GFP* chromosome from *nos-Cas9* and *QUAS-6XGFP*. To make *Hmx-GFP*, pHACK(HmxC)-DONR(sfGFP-2A-QF2) was injected into embryos of *y nos-Cas9.P^ZH-2A^ w* by Rainbow Transgenics. The injected animals were crossed to *QUAS-6XGFP*. GFP-positive animals were selected from their progeny at larval stages and later crossed to the *TM6B* balancer to separate the *Hmx-GFP* chromosome from *nos-Cas9* and *QUAS-6XGFP*. Both strains were confirmed by genomic PCR and sequencing.

### Generation of *Gal4* transcription reporters via the trojan exon method

The *Rbp6-Gal4*, *rdgC-Gal4*, and *yellow-b-Gal4* driver lines were generated by converting specific MiMIC insertions into T2A-Gal4 alleles using the Trojan exon method (*50*). The starting strains were *Mi*(*10*)*Rbp6^MI08679^*, *Mi*(*10*)*rdgC^MI06989^* and *Mi*(*10*)*yellow-b^MI12249^*, respectively. A donor transgene carrying the *SA-T2A-Gal4* cassette [pC-(lox2-attB-SA-T2A-Gal4-Hsp70)] was introduced into the MiMIC backgrounds. Males carrying both the donor and the MiMIC element were crossed to females expressing germline Cre (*Cre(y⁺), vas-int*) to generate founder flies that contained all three genetic components. The founder flies were crossed to *UAS-GFP* reporter lines for subsequent screening of GFP-positive progeny at larval stages. The candidates were recovered, and the converted Gal4 chromosomes were selected by crossing to appropriate balancers stocks. Successful integrations were confirmed via genomic PCR and sequencing.

### Live imaging and microscopy

Live imaging of larval sensory neurons was performed following established protocols (*45*). Larvae were staged and collected at specific developmental time points: 24 h AEL (first instar), 96 h AEL (late third instar), and 120 h AEL (wandering third instar). Animals were mounted in glycerol on glass slides and imaged using either a Leica SP8 or Leica STELLARIS 5 confocal microscope. To visualize whole dendritic arbors, images were acquired using a 20× (NA 0.8) oil immersion objective with the pinhole set to 2.5 Airy units and a z-step size of 1.5 µm. For detailed imaging of the soma and proximal dendrites, a 40× (NA 1.3) oil immersion objective was employed with the pinhole set to 2.0 Airy units and a z-step size of 1.0 µm. For standard morphological quantification, analysis was restricted to da neurons within abdominal hemi-segments A2 through A4. For clonal analysis experiments (e.g., CoinFLP and MAGIC), where clone induction is stochastic, neurons from other abdominal segments were also included to maximize sample size.

### Image analysis and quantification

All image processing and analysis were performed using the Fiji/ImageJ software package. Neural morphology quantification, including measurements of terminal dendrite number and total dendrite length, was conducted using a custom ImageJ plugin previously established by our laboratory (*20*). The plugin is available at Zenodo (doi:10.5281/zenodo.11585494). To extract morphological features, neuronal structures were segmented using the “Auto Local Threshold” algorithm and subsequently skeletonized into single-pixel-wide representations. Skeleton lengths were computed based on pixel distances (*45*). For quantitative analysis, terminal dendrite number was defined as the count of branches with a branch order of 1, and total dendrite length was calculated as the sum of the lengths of all dendritic branches across all orders.

### Mosaic analysis by gRNA-induced crossing-over (MAGIC) for sparse axonal labeling

To visualize the fine anatomical details of individual axon terminals within the ventral nerve cord, we employed the Mosaic Analysis by gRNA-Induced Crossing-over (MAGIC) technique established in our laboratory (*31*). This method induces somatic recombination via CRISPR/nCas9-mediated DNA lesions. Specifically, mosaic clones were induced on chromosome arm 3R using *zk-nCas9(H840A)* in combination with the *gRNA-81F(Gal80)* marker transgene, which ubiquitously expresses the repressor Gal80 and a gRNA targeting the pericentromeric region at 81F (*33, 51*). In neuronal precursors, the nCas9 nickase introduces single-strand nicks near the centromere, stimulating mitotic recombination between homologous chromosomes. Following chromosome segregation, daughter cells that become homozygous for the chromosome lacking the gRNA-marker lose the Gal80 repressor. This stochastic loss leads to the derepression of the pan-da driver *Gal4^21-7^*, which subsequently activates *UAS-CD4-tdTom* and *UAS-Hmx-RNAi* expression, resulting in the sparse, positive mosaic labeling of individual neurons against an unlabeled background.

### Transient expression of transgenes with Gal80^ts^

Temporal regulation of gene expression was achieved using the temperature-sensitive Gal80 (Gal80^ts^) system, where Gal4 activity is repressed at the permissive temperature (18°C) and active at the restrictive temperature (29°C). For all experiments, embryos were collected at 18°C over a 6-hour window. For induction of KD only in early embryonic/larval development (Figure 5A), collected embryos were reared at 29°C for 24 h and subsequently shifted to 18°C for the reminder of the larval development. For induction of KD in later larval development (Figure 5F), embryos were initially grown at 18°C and shifted to 29°C at various developmental stages (equivalent of 0–5 days AEL at 25°C). The first batch of animals in those vials that reached wandering 3^rd^ instar larvae after specific days (0-5 days) were selected for imaging. For expression of human genes, larvae were grown initially at 18°C and then transferred to 29°C. Animals that reached wandering 3^rd^ instar larvae after 72 h (for *CUX1* and *HMX1*) or 96 h (for *HMX2* and *HMX3*) at 29°C were selected for imaging. Unless otherwise noted (e.g., L1 imaging), all phenotypic analyses were performed at the wandering 3^rd^ instar larval stage.

### Larval brain dissection and imaging

Larval brain dissection was performed following a previously established protocol (*52*). Wandering 3rd instar larvae were selected for analysis. Initially, larvae were partially dissected to remove the majority of visceral and epidermal tissues, retaining the brain complex and associated connecting structures. The samples were fixed in 4% formaldehyde in PBS for 15 minutes, followed by two 10-minute washes in 0.2% PBST (PBS containing 0.2% Triton X-100). Fine dissection was subsequently performed in fresh PBS to remove all remaining extraneous tissues and isolate the central nervous system. The processed brains were mounted on glass slides using SlowFade Diamond Antifade Mountant (Thermo Fisher Scientific), and a coverslip was applied with light pressure. The ventral nerve cord (VNC) was imaged using either a Leica SP8 or Leica STELLARIS 5 confocal microscope equipped with a 20× (NA 0.8) oil immersion objective.

### Gentle touch behavior test

Larval mechanosensitivity was evaluated using a calibrated eyelash probe following the established Kernan scoring scale (*12, 53*). Wandering third-instar larvae were acclimated on 2% agar plates for 5 min, then received four gentle strokes across the thoracic and abdominal segments (A1–A6) during active locomotion. Responses were recorded and scored as follows: 0, no response; 1, brief hesitation/slowing; 2, anterior retraction (head withdrawal); 3, retraction followed by a turn of <90°; and 4, retraction followed by a turn of >90° or backward movement. Assays were performed double-blind across three biological replicates (n=30 per genotype), and the cumulative scores (0–16 per larva) were used for statistical analysis.

### Single nuclear RNA-seq (snRNA-seq)

#### Nuclei isolation and preparation

Nuclei isolation was adapted from a previously described protocol (*54*). To ensure RNA integrity, all equipment including Dounce homogenizers was cleaned with 70% ethanol and nuclease-free water, followed by heat sterilization. Dissected larval body walls were collected in ice-cold PBS and pelleted by a brief centrifugation. For each batch of collected larval body walls, the supernatant was removed, and the pellet was snap-frozen in liquid nitrogen and stored at -80°C. For downstream processing, the tissue pellet (fresh or frozen) was resuspended in 1 mL of fresh, ice-cold Homogenization Buffer (250 mM sucrose, 10 mM Tris [pH 8.0], 25 mM KCl, 5 mM MgCl2, 0.1% Triton X-100, 0.5% RNasin Plus, 1× protease inhibitor cocktail, and 0.1 mM DTT). Nuclei were released by Dounce homogenization on ice, performed with 20 strokes of the loose pestle followed by 40 strokes of the tight pestle. The homogenate was filtered through a 35–40 µm cell strainer and centrifuged at 1,000 × g for 10 min at 4°C. The supernatant was carefully discarded, and the nuclear pellet was resuspended in 300–600 µL of Resuspension Buffer (1× PBS supplemented with 0.5% BSA and 0.5% RNasin Plus). The suspension was triturated by pipetting (>20 times) to ensure a single-nucleus suspension and subsequently filtered through a 40 µm Flowmi cell strainer. Prior to sorting, a small aliquot was stained with DAPI to verify nuclear integrity and density under an epifluorescence microscope.

#### Fluorescence-activated cell sorting (FACS)

For sorting, DAPI (Sigma) was added to the nuclear suspension to a final concentration of 1 µg/mL to label all nuclei. Sorting was performed on a BD FACSAria Fusion flow cytometer using a 100 µm nozzle. The gating strategy was hierarchically set as follows: (1) debris removal using FSC-A vs. SSC-A; (2) doublet discrimination using DAPI-A vs. DAPI-H; (3) selection of DAPI-positive nuclei; and (4) enrichment of da neuron nuclei based on EGFP fluorescence intensity (labeled by *Gal4^RluA1^>UAS-Lam-EGFP*). Sorted nuclei were collected into tubes containing resuspension buffer and immediately processed for library construction.

#### SnRNA-seq library preparation and sequencing

Single-nucleus gene expression libraries were constructed using the Chromium Next GEM Single Cell 3’ Reagent Kits v3.1 (10x Genomics) according to the manufacturer’s instructions. Library quality was assessed using an Agilent 2100 Bioanalyzer. Sequencing was performed on an Illumina NextSeq 2000 platform to achieve a depth of approximately 50000 reads per nucleus.

### SnRNA-seq data analysis

Raw sequencing data were demultiplexed, aligned to the *Drosophila melanogaster* reference genome (BDGP6.32), and counted using Cell Ranger (V8.0, 10x Genomics). The resulting gene-barcode matrices were imported into the Seurat R package (v4) for downstream analysis, including quality control, filtering, and clustering.

#### Quality control and filtering

Nuclei were filtered based on the following criteria: number of detected genes (nFeature_RNA > 200 and < 2500) and percentage of mitochondrial reads (< 5%).

#### Dimensionality reduction and clustering

Data were normalized using the LogNormalize method and scaled. Highly variable features were identified using the FindVariableFeatures function. Principal Component Analysis (PCA) was performed, and the top 45 principal components were used for non-linear dimensionality reduction (UMAP) and clustering (FindNeighbors and FindClusters).

#### Marker identification and annotation

Differentially expressed genes (DEGs) were identified using the FindAllMarkers function with a Wilcoxon Rank Sum test. Cell types were annotated based on the expression of canonical markers (*ct*, *Hmx*, *nompC*, *ppk*, etc.). For comparative analysis between genotypes (Control, *ct RNAi*, *Hmx RNAi*), datasets were integrated using Harmony to correct for batch effects before clustering.

### Cross-Species Transcriptomic Comparison

Publicly available human dorsal root ganglion (DRG) single-nucleus RNA-seq datasets were retrieved from (*34*). To enable cross-species alignment, we generated a curated list of 313 high-confidence orthologous genes between *Drosophila melanogaster* and *Homo sapiens*, encompassing ion channels, transcription factors, signaling molecules, and cell adhesion proteins (Table S2). Transcriptional similarity between fly da neuron subtypes and human DRG clusters was quantified using the MetaNeighbor R package. The analysis was restricted to the identified ortholog set to calculate Area Under the Receiver Operating Characteristic (AUROC) scores, which assess the replicability of cell-type identity across datasets based on neighbor voting. The resulting similarity matrix was visualized as a heatmap using the pheatmap package. For gene expression visualization, normalized expression values of selected orthologs were projected onto the pre-computed UMAP embeddings of the respective datasets.

### Statistical analysis

Statistical analyses were performed using the R software. For comparisons between two groups, Student’s t-test was used. For comparisons involving three or more groups, one-way analysis of variance (ANOVA) was applied, followed by Tukey’s post-hoc test to determine significant differences between means. Statistical significance is indicated as *p < 0.05, **p < 0.01 and ***p < 0.001; n.s. denotes not significant.

## SUPPLEMENTARY FIGURES

**Figure S1.**
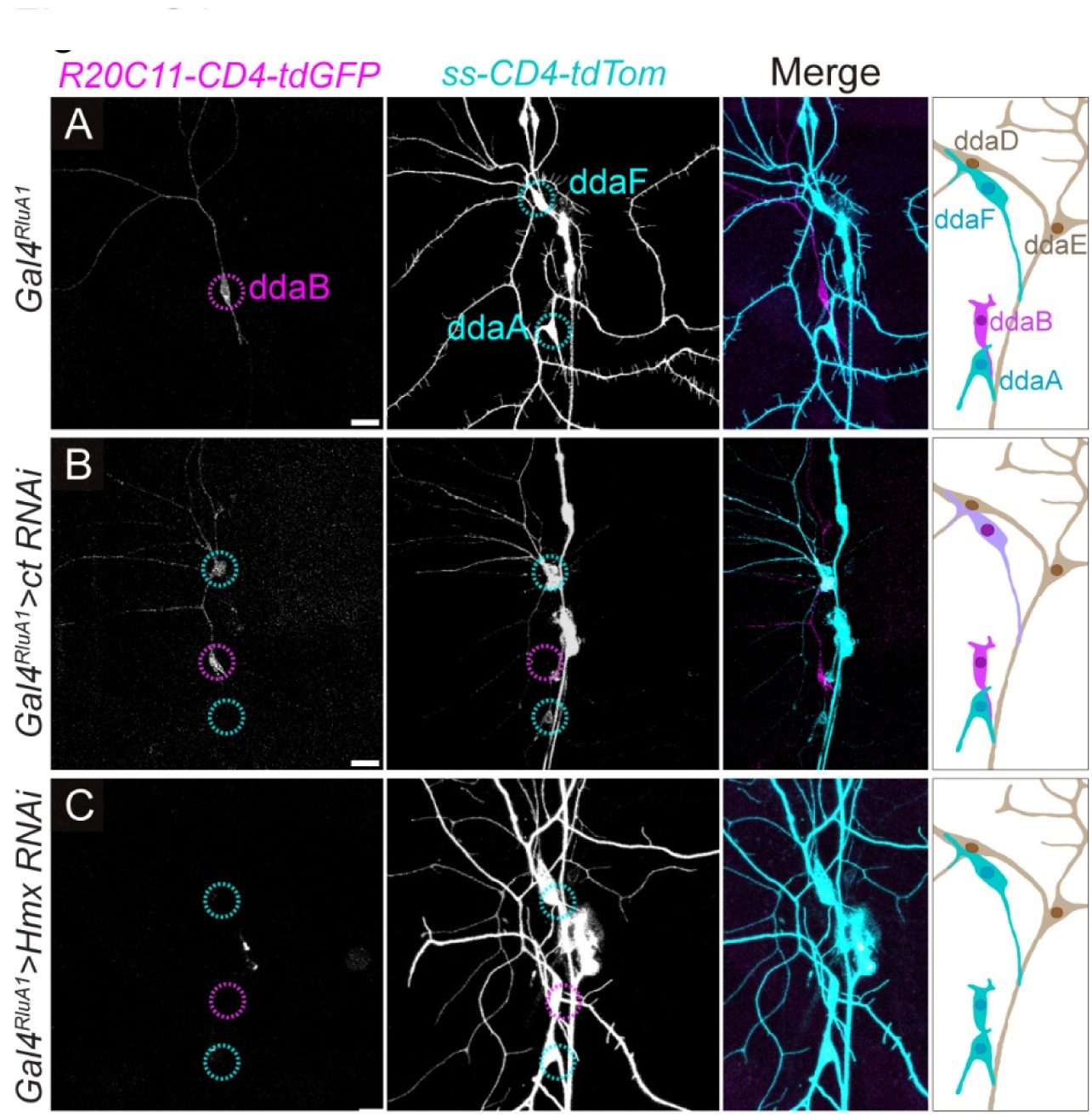
*ct* and *Hmx* KD cause marker switches between C2da/C3da neurons. **(A–C)** Expression of additional markers (*R20C11-CD4-tdGFP* for ddaB and *ss-CD4-tdTom* for ddaA/F) in dorsal da neurons of WT control (**A**), *ct RNAi* (**B**), and *Hmx RNAi* (**C**) in late 3rd instar larvae. Scale bars: 20 μm

**Figure S2.**
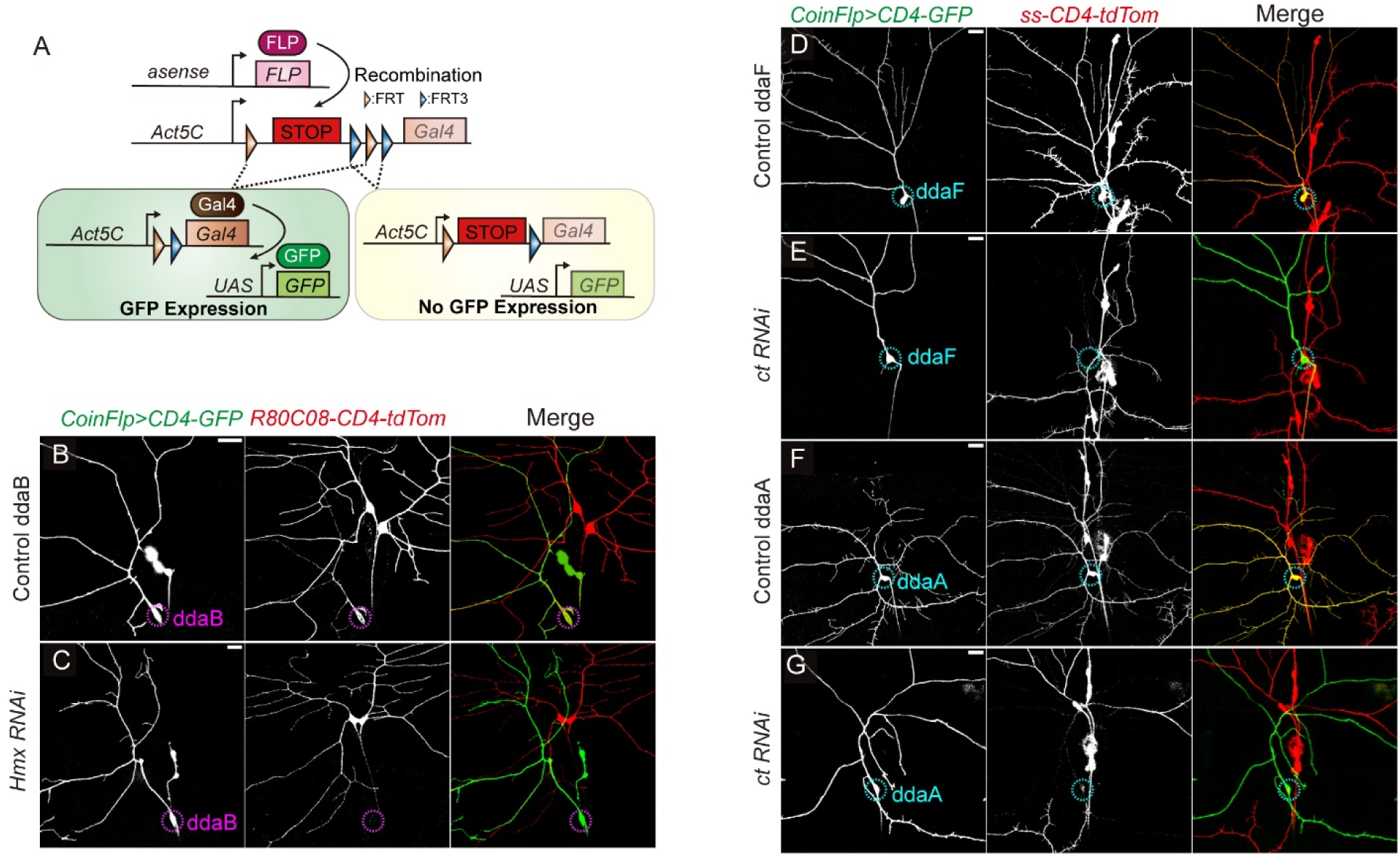
*ct* and *Hmx* manipulations in single neurons using the CoinFlp system. **(A)** Schematic illustration of the CoinFLP system used to label single-neuron clones. *asense-FLP* induces stochastic recombination of one of two incompatible sets of FRT sites in neural progenitors. Successful excision of the STOP cassette in a small subset of neuronal lineages triggers constitutive *Act5C*-driven Gal4 expression, resulting in labeling of individual neuronal clones by *UAS-CD4-GFP*. This allows for high-resolution morphological analysis of single clones against a background of unlabeled cells. **(B–G)** Single-neuron clones generated by CoinFLP. *CoinFlp>CD4-tdGFP* indicates the clones. The identities of the clones were determined based on overlap with (in WT clones) or the loss of *R80C08-CD4-tdTom* (C1da/C2da marker) and *ss-CD4-tdTom* (C3da marker) in *Hmx* and *ct* KD clones, respectively. Scale bars: 20 μm.

**Figure S3.**
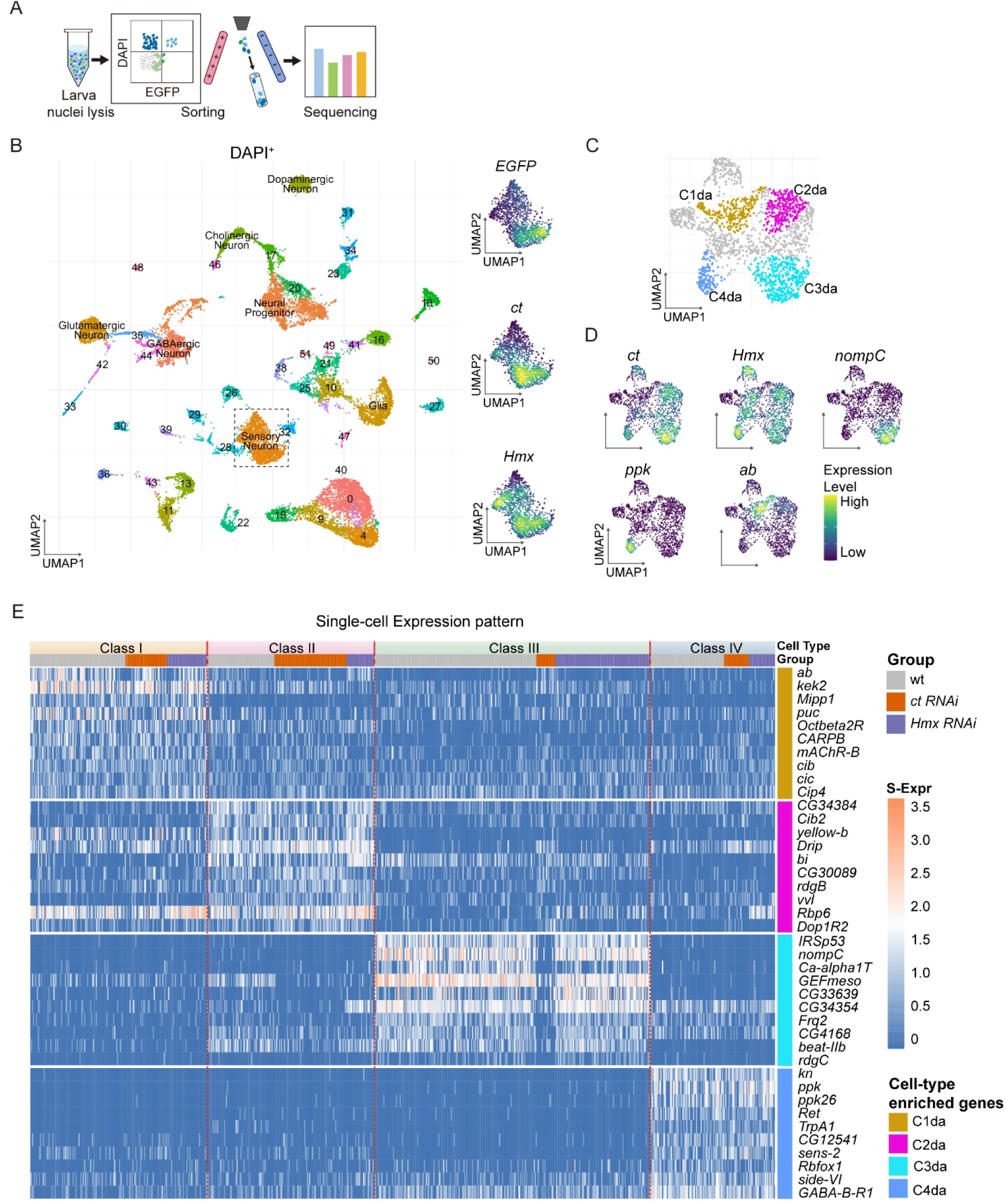
snRNA-seq analyses of da neurons. **(A)** Scheme of snRNA-seq workflow. Nuclei were isolated from larval body walls of *Gal4^RluA1^>UAS-Lam-EGFP* and collected by fluorescence-activated cell sorting (FACS) based on DAPI (nuclei) and EGFP (da neurons) signals prior to sequencing. **(B)** UMAP visualization of all sequenced nuclei. Some clusters are annotated based on transcriptional signatures. Feature plots (right) showing expression of *EGFP*, *ct*, and *Hmx* were used to identify the sensory neuron cluster (dashed box). **(C)** Re-clustering of the sensory neuron cluster (dashed box in **B**). Da neuron subtypes are color-coded. **(d)** Expression of *ct*, *Hmx*, and known cell-type markers on the UMAP from (**C**). *nompC* marks C3da; *ppk* marks C4da; *ab* marks C1da. Color intensity represents expression level. **(E)** Single cell heatmap showing expression of top enriched genes for each da neuron class. Columns represent individual cells grouped by cluster assignment (Class I–IV) and genotype. Rows are organized by cell-type specificity (C1da to C4da, top to bottom). Color intensity indicates scaled expression level (S-Expr).

**Figure S4.**
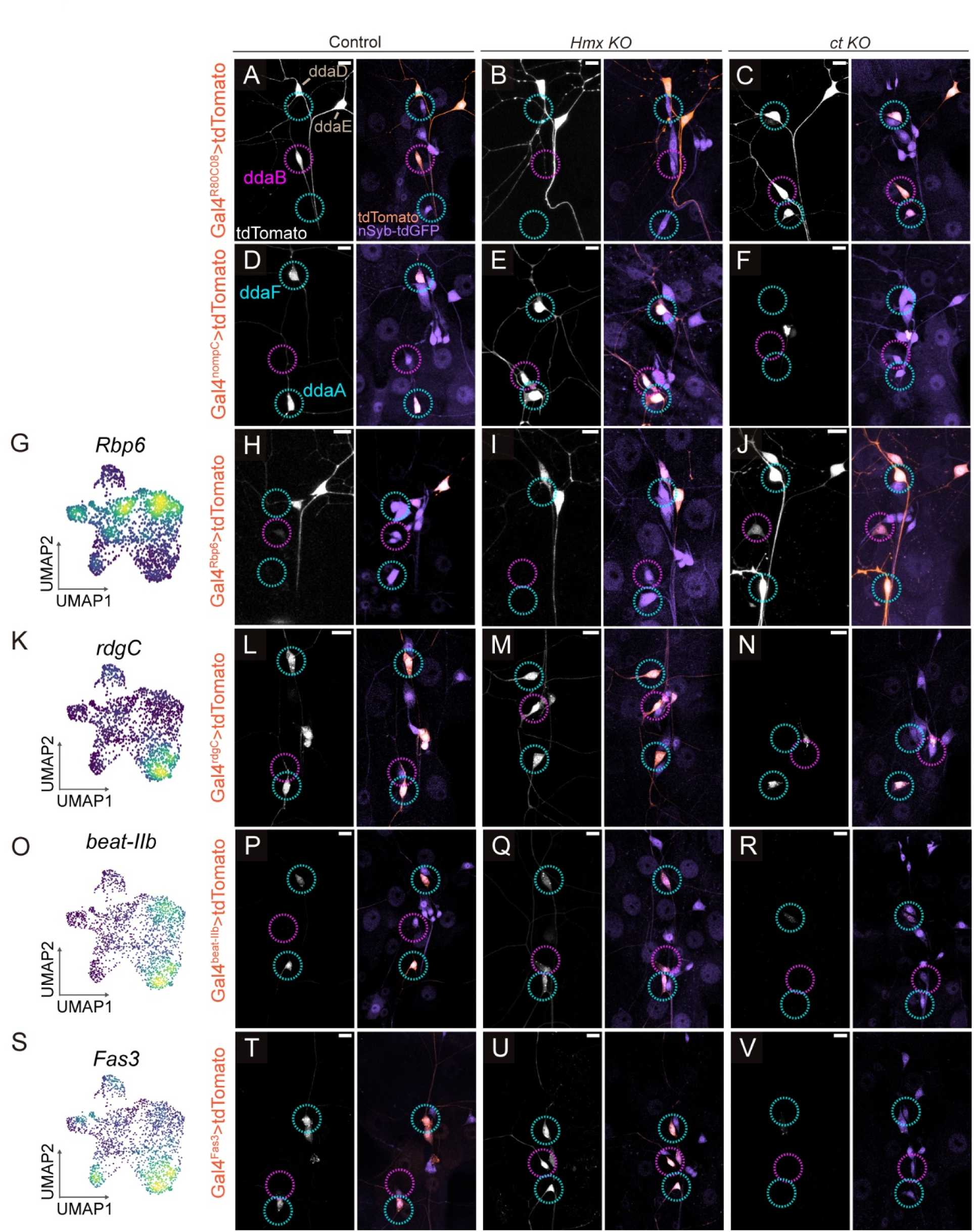
*In vivo* validation of sensory effector reprogramming and cell-type conversions following *ct* and *Hmx* knockout. **(A–F)** Validation of *ct* and *Hmx* KO by *ase-Cas9* using C2da-specific *Gal4^R80C08^* (**A-C**) and C3da-specific *Gal4^nompC^*(**D-F**) in the WT control. **(G-V)** Validation of additional DEGs with corresponding Gal4 reporters. UMAP plots of C2da-enriched *Rbp6* (**G**) and C3da-enriched genes (*rdgC*, **K**; *beat-IIb*, **O**; *Fas3*, **S**) serve as single-cell expression references. The respective Gal4 lines drive *UAS-tdTomato* expression in Control (**H, L, P, T**), *Hmx KO* (**I, M, Q, U**), and *ct KO* (**J, N, R, V**) larvae. The pan-neuronal marker *nSyb-tdGFP* was included to facilitate identification of neurons. Magenta circles indicate C2da (ddaB) neurons, and cyan circles indicate C3da (ddaA/ddaF) neurons. Scale bars: 10 μm.

**Figure S5.**
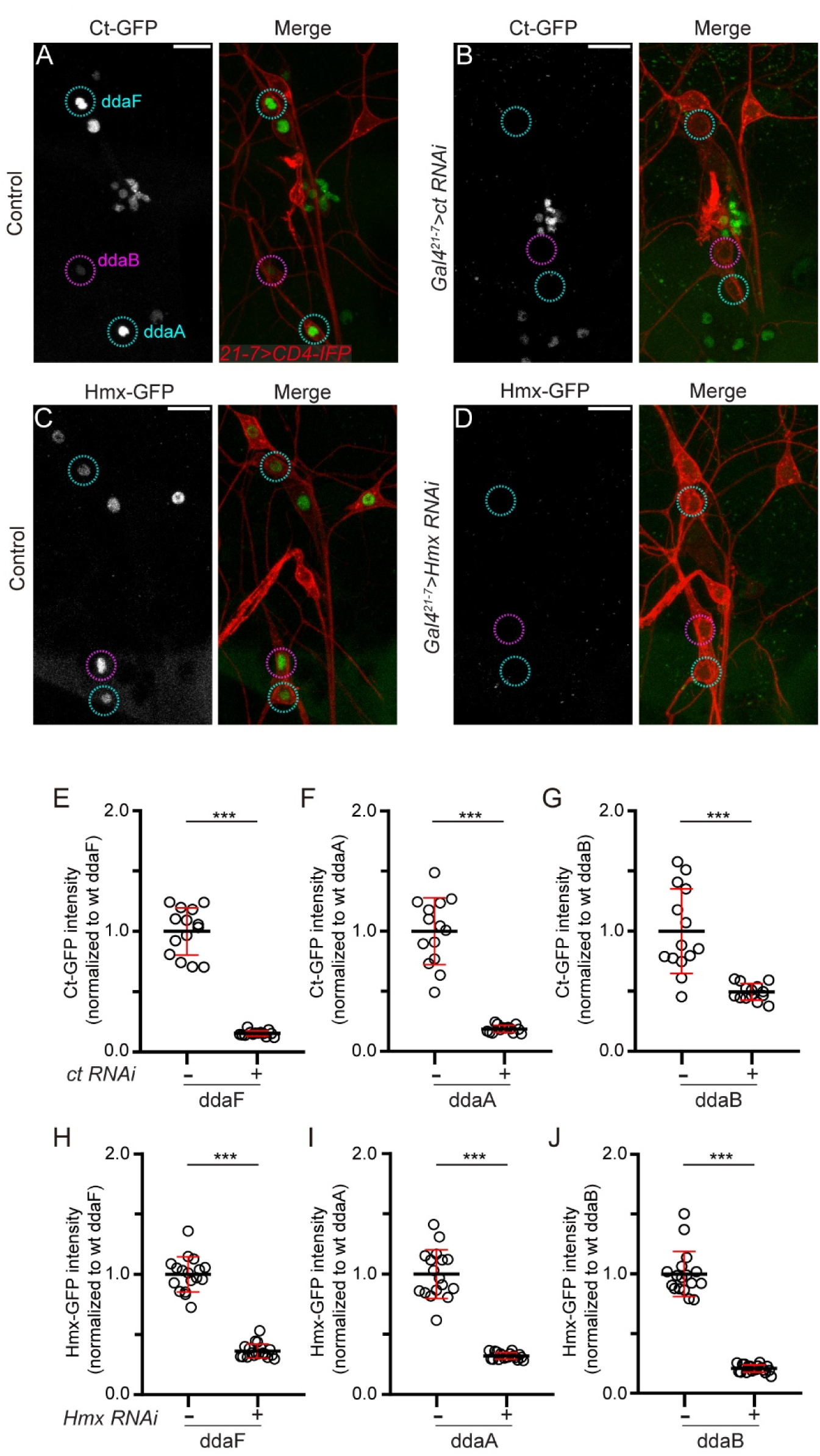
Validation of *ct* and *Hmx* KD efficiency. **(A–D)** Expression of endogenous *Ct-GFP* **(A, B)** and *Hmx-GFP* **(C, D)** in da neurons of control **(A, C)**, *ct* RNAi **(B)**, and *Hmx* RNAi **(D)** larvae. RNAi expression was driven by *Gal4^21-7^*. Da neurons were visualized by *UAS-CD4-IFP* expression. Cyan circles indicate C3da neurons (ddaA/F); magenta circles indicate C2da neurons (ddaB). Scale bars: 10 μm. **(E–J)** GFP fluorescence intensity of *Ct-GFP* **(E–G)** and *Hmx-GFP* **(H–J)** in ddaF (**E**, **H**), ddaA (**F**, **I**), and ddaB (**G**, **J**) neurons. Values are normalized by the mean intensity of control neurons of the same type. Sample sizes (n = neurons): n = 12–18 per genotype. Unpaired t-test (***p<0.001). Black bars, mean; red bars, SD.

**Figure S6.**
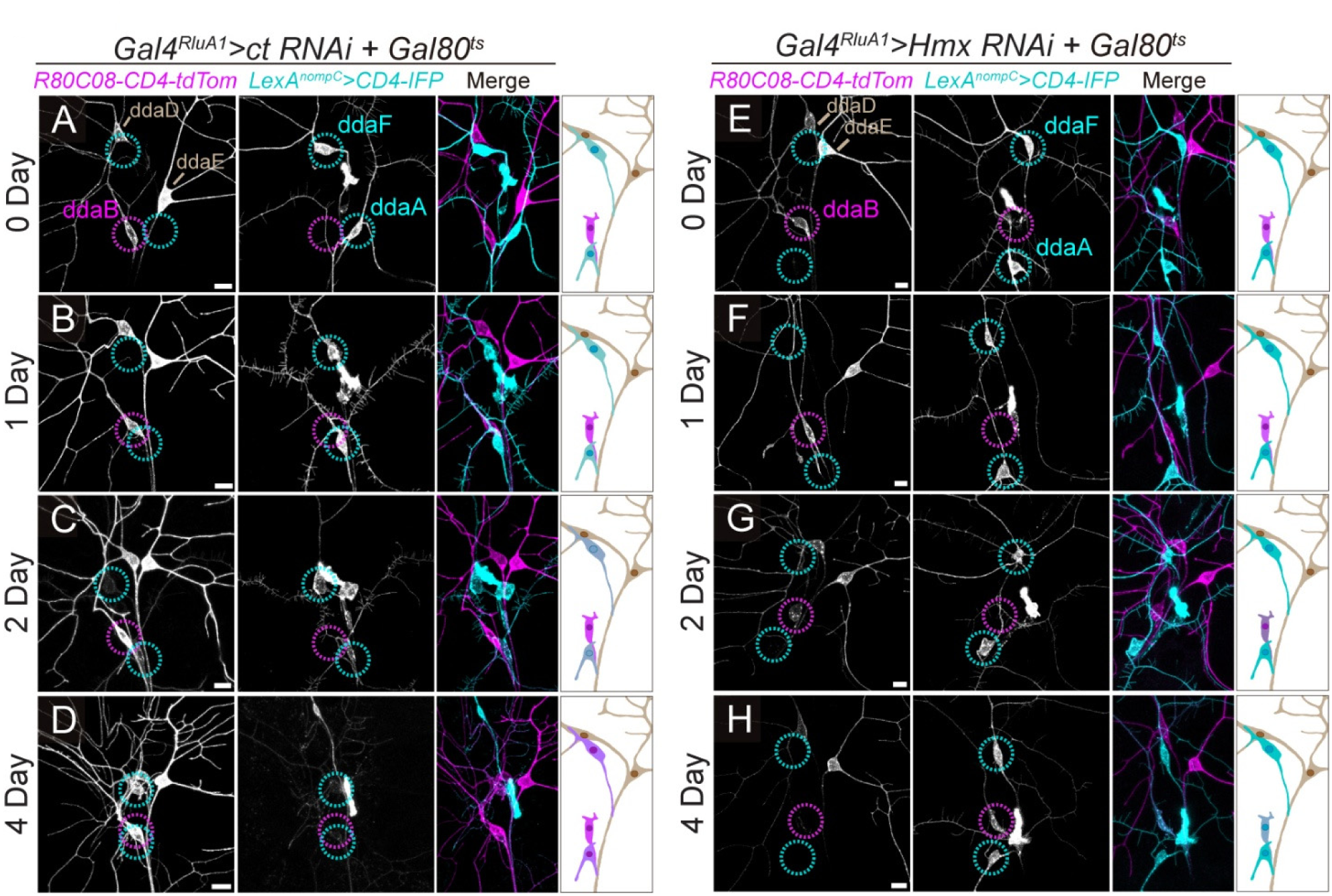
Temporal profiles of cumulative transformation. **(A-H)** Expression of markers (*R80C08-CD4-tdTom* for C1da/C2da and *LexA^nompC^>CD4-IFP* for C3da) in da neurons following 0-day, 1-day, 2-day, and 4-day induction of *ct RNAi* (**A-D**) or *Hmx RNAi* (**E-H**). RNAi expression is driven by *Gal4^RluA1^* under *Gal80^ts^* control (29°C for induction). Cyan circles indicate C3da neurons (ddaA/F); magenta circles indicate C2da neurons (ddaB). Diagrams on the right illustrate cell conversion outcomes. In all image panels, scale bars, 10 μm.

**Figure S7.**
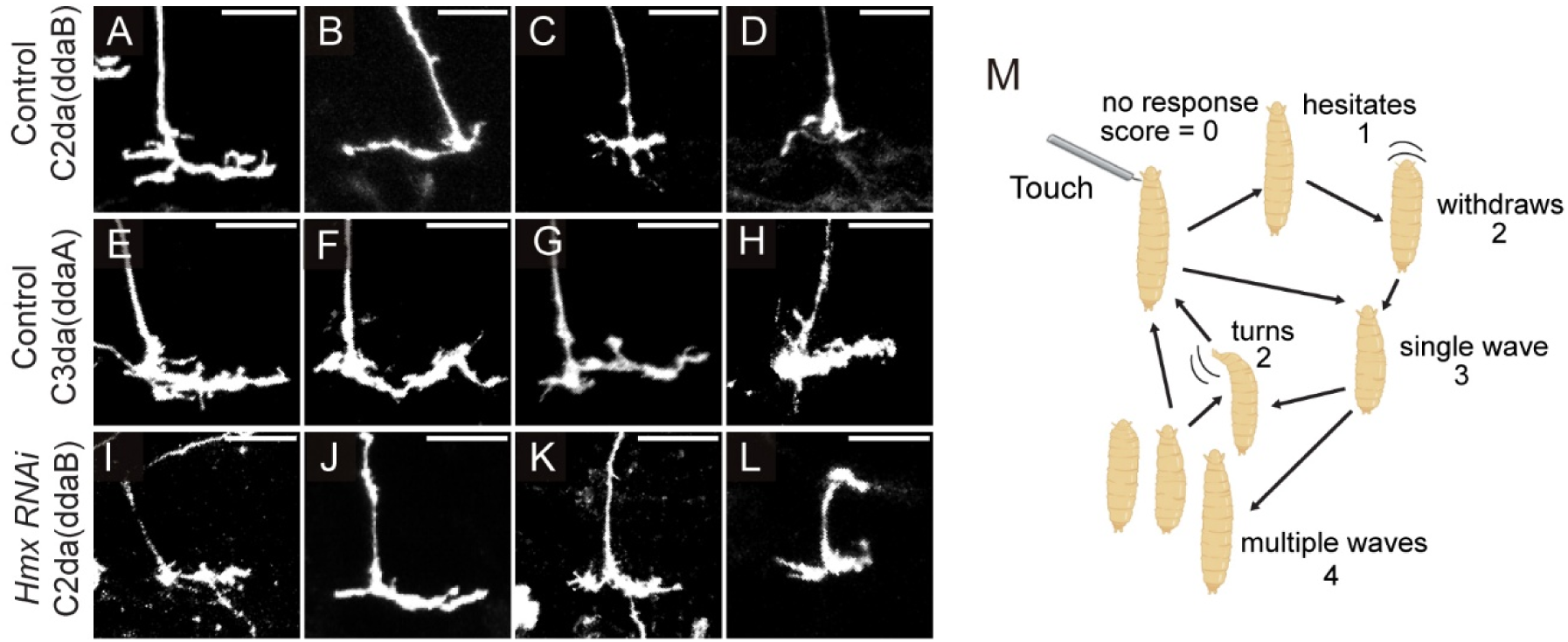
Additional examples of axon projections and behavioral assay scoring scheme. (**A–L**) Additional examples of axon terminal morphologies of WT ddaB (C2da) (**A–D**), WT ddaA (C3da) (**E–H**), and *Hmx RNAi* ddaB (**I–L**) clones generated by the MAGIC technique. (**M**) Diagram of the gentle touch assay and the scoring scheme (0–4), adapted from (*12*).

**Figure S8.**
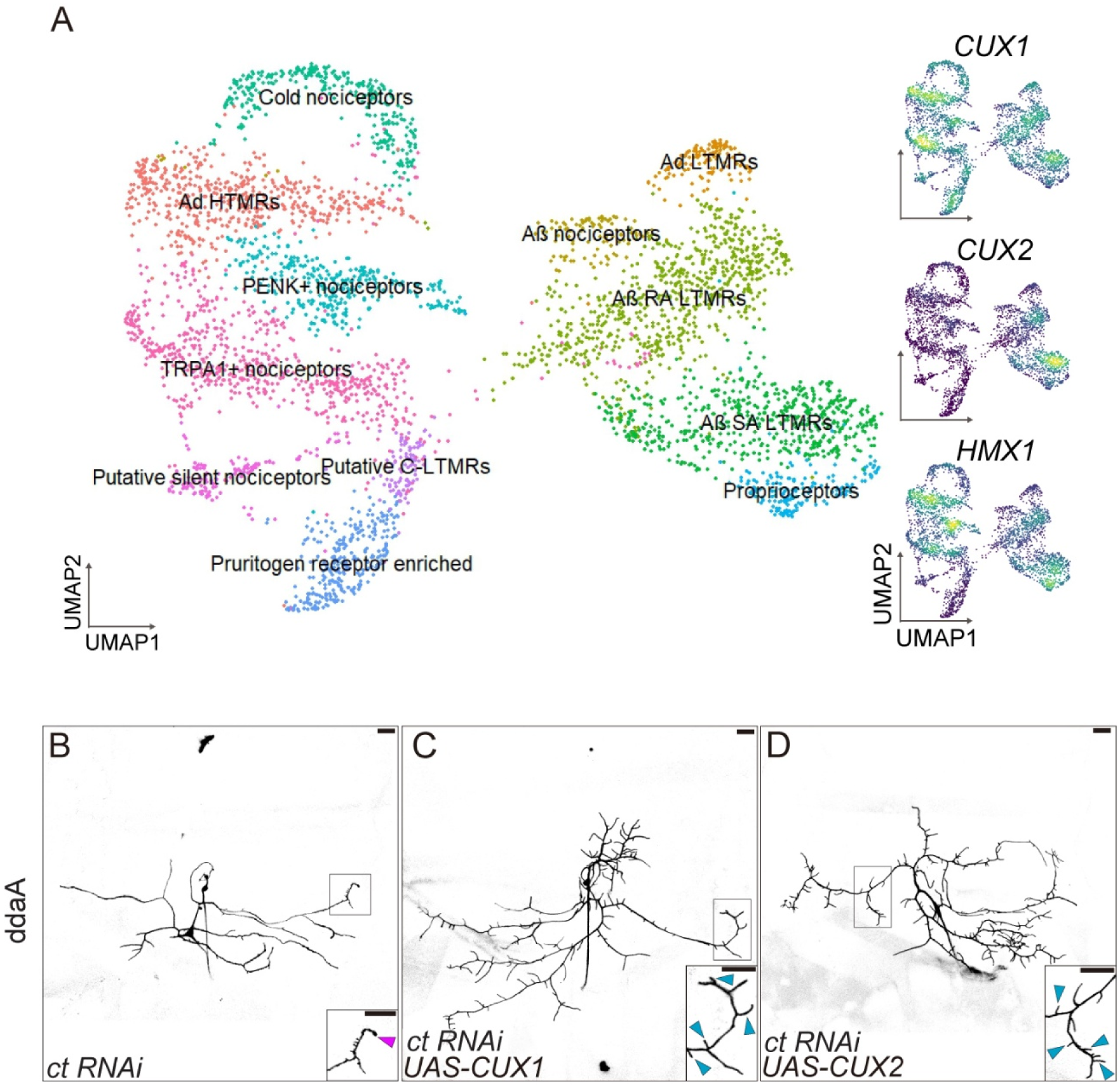
Similarities of somatosensory transcriptional signatures between *Drosophila* da neurons and human DRG neurons. **(A**) UMAP of human DRG neurons based on the dataset and classification from (*34*). Expression of CUX1, CUX2, and HMX1 is shown on the right. **(B–D)** CoinFlp clones of ddaA neurons expressing *ct* RNAi alone **(B)**, or co-expressing *UAS-CUX1* **(C)** or *UAS-CUX2* **(D)**. Insets show high-magnification views of dendritic terminals. Cyan arrowheads indicate dendritic spikes; magenta arrowheads indicate varicosities. Scale bars: 25 μm.

## Notes

### Competing Interest Statement

The authors have declared no competing interest.

